# Transient systemic inflammation in adult male mice results in underweight progeny

**DOI:** 10.1101/2020.10.30.358697

**Authors:** Sushama Rokade, Manoj Upadhya, Dattatray S. Bhat, Nishikant Subhedar, Chittaranjan S. Yajnik, Aurnab Ghose, Satyajit Rath, Vineeta Bal

**Author notes:** These authors contributed equally (Satyajit Rath, Vineeta Bal). Name and postal address of corresponding author*: Vineeta Bal, Department of Biology, Indian Institute of Science Education and Research, Homi Bhabha Road, Pune 411008, India.).

## Abstract

**Problem:** While the testes represent an *immune privileged* organ, there is evidence that systemic inflammation is accompanied by local inflammatory responses. We therefore examined if transient systemic inflammation caused any inflammatory and functional consequences in murine testes.

**Method of Study:** Using a single systemic administration of Toll-like receptor (TLR) agonists [lipopolysaccharide (LPS) or peptidoglycan (PG) or polyinosinic-polycytidylic acid (polyIC)] in young adult male mice, we assessed testicular immune-inflammatory landscape and reproductive functionality.

**Results:** Our findings demonstrated a significant induction of testicular TNF-α, IL-1β and IL-6 transcripts within 24 h of TLR agonist injection. By day 6 these cytokine levels returned to baseline. While there was no change in caudal sperm counts at early time points, eight weeks later, two-fold decrease of sperm count and reduced testicular testosterone levels was evident. When these mice were subjected to mating studies, no differences in mating efficiencies or litter sizes were observed compared to controls. Nonetheless, the neonatal weights of progeny from LPS/PG/polyIC treated sires were significantly lower than controls. Postnatal weight gain up to three weeks was also slower in the progeny of LPS/polyIC treated sires. Placental weights at 17.5 days post-coitum were significantly lower in females mated to LPS and polyIC treated males. Given this likelihood of an epigenetic effect, we found lower testicular levels of histone methyl-transferase enzyme, mixed-lineage leukemia-1, in mice given LPS/PG/polyIC eight weeks earlier.

**Conclusion:** Exposure to transient systemic inflammation leads to transient local inflammation in the testes, with persistent sperm-mediated consequences for fetal development.

## Introduction

As the primary site for the production of spermatozoa and testosterone, mammalian testes are crucial determinants of male fertility (1). Further, unlike oogenesis in the female, testicular spermatogenesis is a continuous process (2). As a result, changes in the testicular tissue microenvironment carry a significant potential for altering spermatozoal programming, with trans-generational consequences (3). The blood-testis barrier and the resulting immune-privileged status of the testes become interesting in this context as well, although they are more commonly invoked from the perspective of prevention of auto-immune responses (4, 5).

Testicular inflammatory responses, both auto-reactive and anti-microbial, particularly anti-viral, are well documented factors in male infertility (6). A number of viruses show testicular tropism, and initiate anti-viral immune responses in the testis leading to inflammation, tissue damage and impaired testosterone and sperm production (7, 8, 9, 10). Systemic inflammatory states mediated by microbial ligands for pattern recognition receptors (PRRs) are known to induce testicular responses as well, although many studies connect such local responses to the maintenance of immune privilege (11, 12, 13). Testicular cells express a broad array of PRRs such as Toll-like receptors (TLRs) and RIG-I-like receptors (14, 15, 16, 17). Using animal models of systemic administration of lipopolysaccharide (LPS), a cell wall component of Gram-negative bacteria, several studies have reported local testicular production of pro-inflammatory cytokines such as TNFα, IL1β, IL6 as well as of reactive oxygen species. Such inflammatory responses have been associated with short-term detrimental effects on testicular functions of spermatogenesis and steroidogenesis, breach of blood-testis barrier, induction of germ cell apoptosis and loss of sperm motility leading to infertility (18, 19, 20, 21).

However, there have been few studies examining if transient systemic inflammation, that induces a similarly transient response in the testes, has any long-term functional consequences on reproductive potential. It has been reported that a single dose of systemic LPS administration can cause damage lasting up to five weeks to the testicular structure in mice with significant decrease in spermatogenesis, meiotic index and epithelial height (22). Another recent study has similarly reported month-long testicular damage upon exposure to a single systemic LPS dose, with apoptosis of spermatogonial germ cells, severely impaired spermatogenesis, thickening of smooth muscle layers, infiltration of immune cells in the interstitial spaces and presence of germ cells in the lumen of cauda epididymis (23). Such effects would be particularly relevant in communities with high microbial exposure loads, where mild to moderate systemic inflammation, commonly asymptomatic or only mildly symptomatic, would be a matter of common occurrence.

On this background, we report an analysis of the long-term functional consequences of transient systemic inflammation in the mouse testes. We report that a single systemic exposure of young adult male mice to TLR ligands showed transient local induction of pro-inflammatory cytokines, followed by a return to baseline within six days. However, sperm counts showed persistent mild reduction. When these male mice were used in mating studies, while they show unimpaired fertility, their progeny were born with significantly lower birth weights and showed persistent post-natal delays in weight gain. This functional epigenetic effect was accompanied by persistent reduction in the testicular levels of a histone methyl-transferase, MLL1. These data open up mechanistic possibilities for inflammation-mediated epigenetic changes in the spermatozoal differentiation programme. Further, they open up translational possibilities as a factor for explaining the persistently high frequency of India’s low birth weight problem.

## Materials and methods

### Animals

C57BL/6 male mice (aged 8 weeks) and C57BL/6 female mice with proven fertility (aged 12-14 weeks) were obtained from the National Facility for Gene Function in Health and Disease (NFGFHD), IISER, Pune. The study was reviewed and approved by the Institutional Animal Ethics Committee (IISER/IAEC/2018-01/05), and was conducted in accordance with Committee for the Purpose of Control and Supervision of Experiments on Animals (CPCSEA)-approved guidelines. Animals were maintained under controlled conditions of temperature and humidity with 12 h light and 12 h dark daily cycle, and were fed *ad libitum*.

For inducing systemic inflammation, 8 week-old male mice were given TLR ligands namely, LPS (5 mg/kg; from *Salmonella typhosa*, Sigma, St. Louis, MO, USA), or PG (5 mg/kg; from *Staphylococcus aureus*, Sigma), or polyIC (10 mg/kg; Sigma) by intra-peritoneal (i.p.) injection. Control mice were similarly given normal saline. Animals were euthanised at 24, 72 and 144 h, or 8 weeks later. Body weights were recorded at each time point prior to euthanasia. Mice were anaesthetised with isoflurane and blood samples were collected via the retro-orbital sinus for serum separation. Subsequently, cardiac perfusion was carried out with ice-cold PBS. Testes, spleen, and brain tissues of perfused animals were collected in TRIsoln reagent (GeNei, Bangalore, India) for RNA extraction and cytokine analysis while caudae epididymides were collected in PBS for measuring the sperm count. Testes from the 8-week groups were frozen in liquid nitrogen for testosterone assays.

### RNA isolation and quantitative reverse transcriptase-PCR (qRTPCR) analysis

Total RNA extracted as above was treated with DNase I (Thermo Scientific, Rockford, USA) to eliminate any potential genomic DNA contamination. The yield and quality of the resulting RNA was evaluated by determining the A260:A280 ratio. RNA (700 ng) was reverse transcribed using iScript cDNA synthesis kit (Biorad, Hercules, CA, USA). The cDNA obtained was used for subsequent real-time quantitative qPCR reactions (CFX96 Thermal Cycler, Bio-Rad). Reaction specificities were validated by melt curve analyses. For all experiments, 18s mRNA was used as internal control. Primer sequences used for the amplification of cDNAs were designed using NCBI Primer BLAST Software (Supplementary Data, Table S1).

### Sperm count

Caudae epididymides from euthanised mice were collected in PBS. Incisions were made in the caudae to release the sperm, which were allowed to exude at 37°C for 30 min. The resulting sperm suspension was washed, re-suspended and counted using a haemocytometer. Sperm counts were performed in replicates for each sample at two dilutions.

### Testosterone assay

Frozen testes were homogenised in 1 ml of ice-cold 1% SDS containing 0.5N NaOH and incubated at 40°C for 3 h on a rotary shaker at 100 rpm. The homogenate was centrifuged at 13000 g for 20 min at 4°C. The resulting testicular supernatants and serum samples were appropriately diluted and testosterone levels were analysed using a testosterone-specific competitive ELISA kit according to manufacturer’s protocol (Diagnostic Biochem Canada Inc, Dorchester, ON, Canada).

### Fertility studies

To study the effect of transient systemic inflammatory stress on fertility, 8 week-old male mice were given a single injection of LPS, PG or polyIC (or PBS as a control) as above. Eight weeks later, the reproductive capacity of these mice was evaluated by cohabitation with healthy syngeneic females. Briefly, each male was caged individually with two females of proven fertility for a maximum period of 20 days. Fertility assessment was done using the proportions of females achieving pregnancy, duration between the beginning of mating and delivery of the pups, and numbers of offspring per litter. Progeny evaluation was done using pup weights from birth up to 3 weeks of age. At 8 weeks of age, blood glucose levels were determined, and progeny mice were euthanised to determine lengths and lean weights.

For assessing placental weights, mating studies were set up as described above, and females were examined daily for the presence of vaginal plugs to indicate successful mating. The day on which vaginal plugs were observed was considered as 0.5 dpc (day post coitum). At 17.5 dpc, pregnant dams were euthanised and placental and fetal weights recorded.

### Serum glucose assays

Blood was drawn retro-orbitally and sera were separated. Glucose levels were estimated using an automated glucose analyser (Dialab, GmbH, A-2351, Wiener Neudorf, Austria).

### Statistical analyses

Data were analysed using Prism software (Graph Pad, San Diego, CA, USA). Student’s t-test was used to determine p values, and p < 0.05 was considered statistically significant.

## Results

### A single systemic exposure to LPS induced transient local testicular inflammatory responses and dysfunction

To test the testicular consequences of transient systemic inflammation, young male mice were given a single i.p. dose of LPS (20), which is a component of the outer membrane of Gram-negative bacteria and a TLR4 agonist. Within 24 h, the mice showed symptoms of ruffled body hair, reduced locomotor and exploration activity (Supplementary Data; Figs. S1-S2) and decreased body weight (Fig. 1A). In concordance with these findings, local expression of a number of inflammation-related cytokine genes such as TNFα, IL1β and IL6 was detectable, not only in the spleen (Fig. S3), but also in immune-privileged sites such as the testes (Figs. 1B-1F) and the brain (Fig. S4). However, these effects were transient; they began to decline by 72 h, and by 144 h after LPS administration, most parameters were essentially back to baseline levels (Fig. 1; Supplementary Data, Figs. S3-S4).

**Figure 1.**
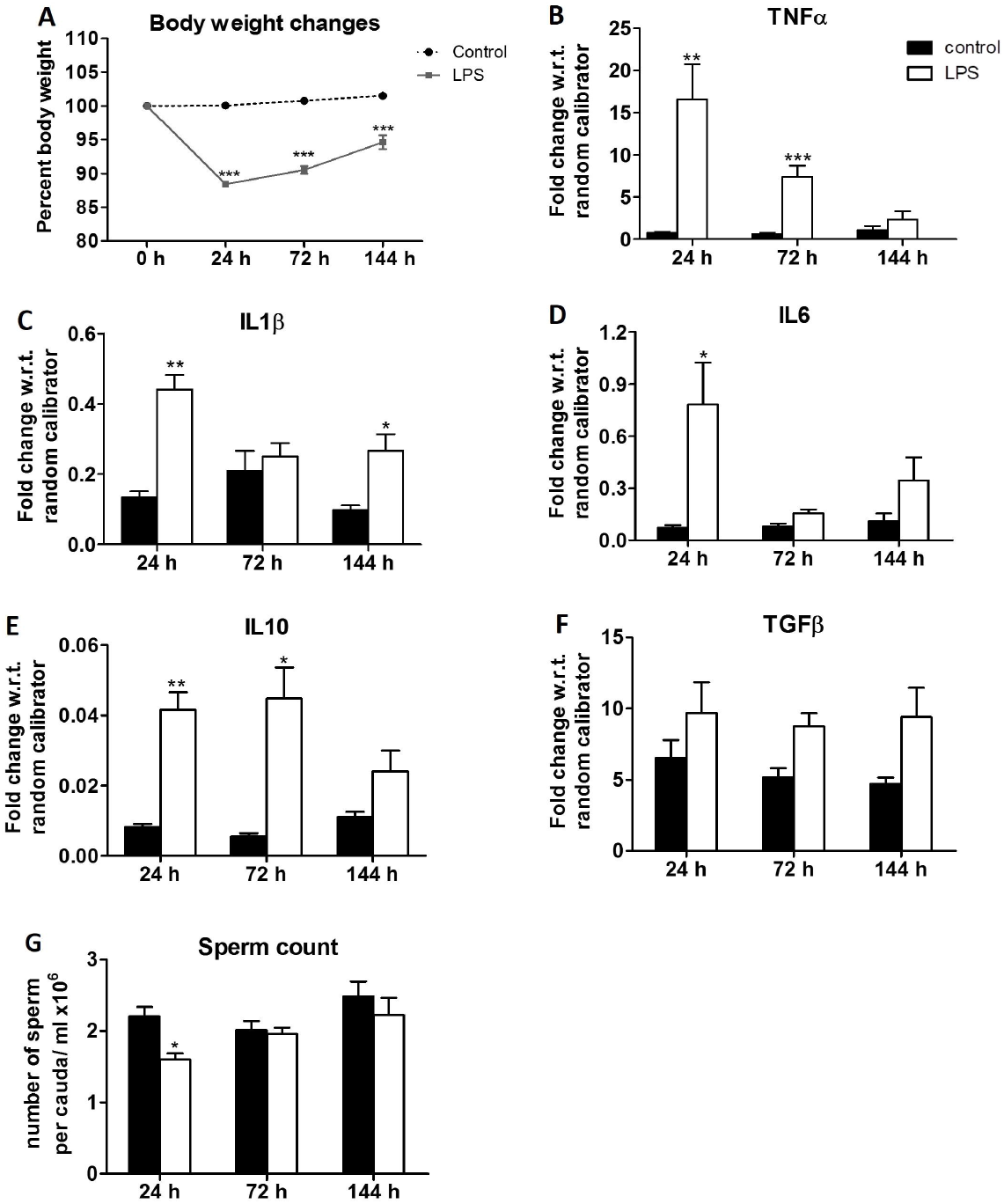
Effects of systemic lipopolysachharide (LPS) injection on body weight, testicular cytokine milieu and caudal sperm counts in mice. Male mice were treated with LPS as described, and (A) body weight changes are shown as normalized to the weight at the beginning of the experiment (n = 9-31) and (B-F) testicular cytokine gene mRNA levels (n = 5), were assayed at various times as shown. Levels of mRNA for TNFα (B), IL1β (C), IL6 (D), IL10 (E) and TGFβ (F) were measured by qRTPCR and normalised values as shown were calculated. (G) Caudal sperm counts at indicated times post-LPS injection (n = 5-11). Data are shown as mean ± s.e.. * p< 0.05, **p<0.01, ***p<0.0001.

In association with these transient effects, the caudal sperm counts were significantly reduced in LPS-treated mice at 24 h. However, by 72 h, sperm counts had returned to normal levels in these mice (Fig. 1G).

### Transient systemic exposure to TLR2 and TLR3 ligands, PG and polyIC, induced a milder testicular inflammation phenotype than that induced by LPS

Since transient exposure to a TLR4 ligand, LPS, induced short-lived local inflammation in the mouse testes, we next examined if ligands for other TLRs also induced similar alterations. Young male mice were given single i.p. doses of either the TLR2 agonist, PG, or the TLR3 agonist, polyIC. Unlike LPS, neither PG nor polyIC induced any transient behavioural alterations (Supplementary Data; Figs. S5-S8). PG induced no transient weight loss either, although polyIC did (Figs. 2A, 3A). However, as seen with the spleen (Fig. S3) PG induced transient local testicular expression of some, though not all, inflammation-related cytokine genes tested (Figs. 2B-2F). While polyIC challenge did not induce any inflammatory response in the spleen within 24 h (Fig. S3), testis showed an induction of IL1β, IL6, IFNγ and IL10 transcripts (Figs. 3B-3G). However, no such expression was reliably detected in the brains of these mice (Supplementary Data; Figs. S9-S10). However, as with LPS treatment, all the detectable effects were transient; by 144 h after TLR ligand administration, most parameters were essentially back to baseline levels (Figs. 2, 3). Unlike after LPS treatment, the caudal sperm counts were unaltered at 24-144 h in PG/ poly IC-treated mice (Figs. 2G, 3H).

**Figure 2.**
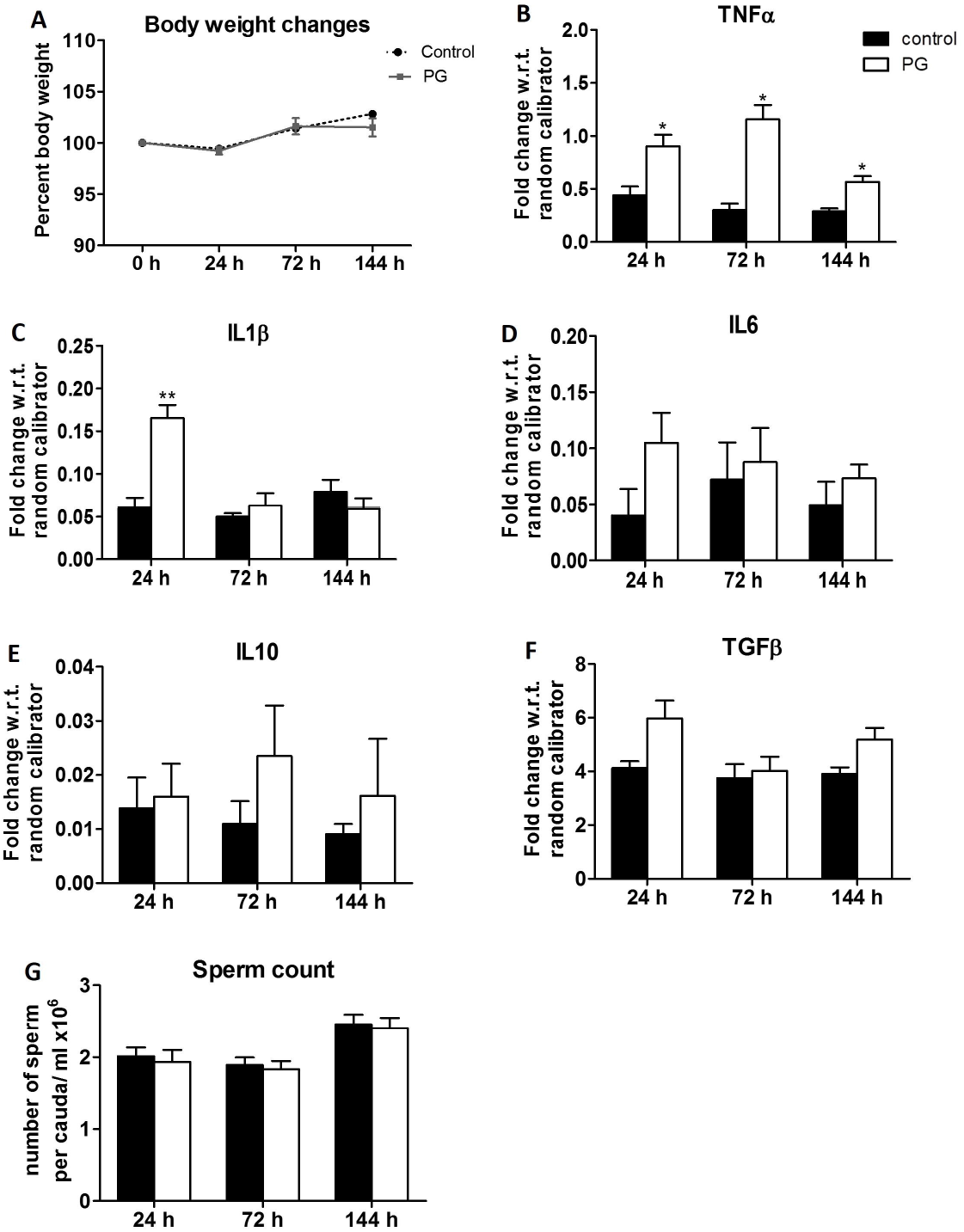
Effects of systemic peptidoglycan (PG) injection on body weight, testicular cytokine milieu and caudal sperm counts in mice. Male mice were treated with PG as described, and (A) body weight changes are shown as normalized to the weight at the beginning of the experiment (n = 5-10) and (B-F) testicular cytokine gene mRNA levels (n = 5), were assayed at various times as shown. Levels of mRNA for TNFα (B), IL1β (C), IL6 (D), IL10 (E) and TGFβ (F) were measured by qRTPCR and normalised values as shown were calculated. (G) Caudal sperm counts at indicated times post-PG injection (n = 5-11). Data are shown as mean ± s.e.. * p< 0.05, **p<0.01.

**Figure 3.**
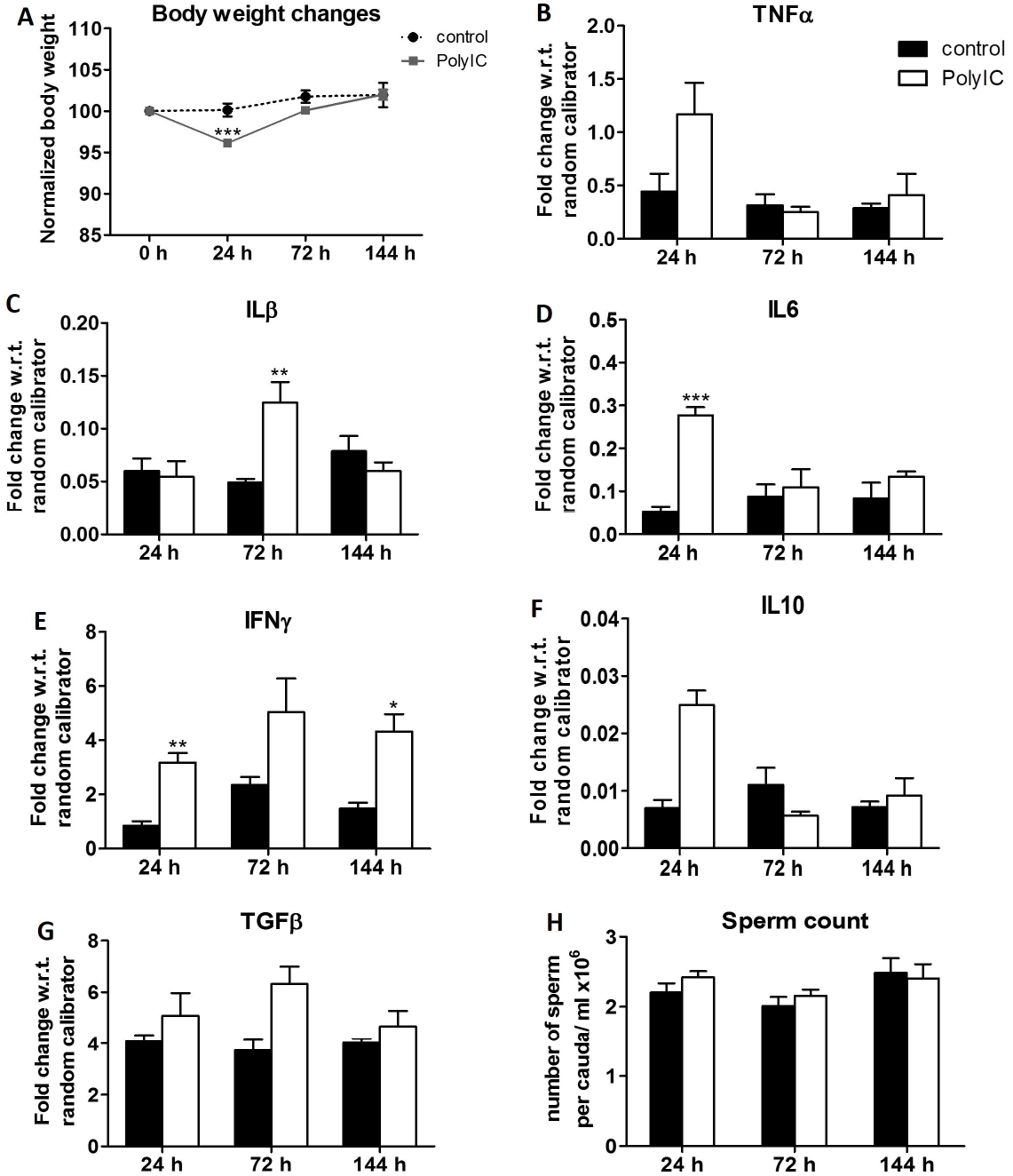
Effect of systemic polyinosinic-polycytidylic acid (polyIC) injection on body weight, testicular cytokine milieu and caudal sperm count in mice. Male mice were treated with polyIC as described, and (A) body weight changes are shown as normalized to the weight at the beginning of the experiment (n = 5-13) and (B-G) testicular cytokine gene mRNA levels (n = 5), were assayed at various times as shown. Levels of mRNA for TNFα (B), IL1β (C), IL6 (D), IFNγ (E), IL10 (F) and TGFβ (G) were measured by qRTPCR and normalised values as shown were calculated. (H) Caudal sperm counts at indicated times post-polyIC injection (n = 5-11). Data are shown as mean ± s.e.. * p< 0.05, **p<0.01, ***p<0.0001.

### Testicular dysfunction at 8 weeks post-LPS/PG/polyIC exposure

It was evident from the data above that challenging young adult mice with a single systemic LPS dose induced only transient inflammatory responses in the testes, which were resolved by 144 h. We next tested if this LPS-mediated transient local testicular inflammation had any long-term functional consequences in the testes. Since the early inflammatory phenotype was milder in mice given PG or polyIC, we used those ligands to test if long-term consequences, if any, were similarly milder.

Interestingly, LPS-exposed male mice showed a two-fold reduction in caudal sperm counts 8 weeks after LPS exposure, and this was true in PG/polyIC-treated mice as well (Fig. 4A). Intra-testicular testosterone levels were significantly lower in LPS- and polyIC-treated mice, but not in PG-treated mice (Fig. 4B). While serum testosterone showed a tendency to lower levels, the differences were not significant (Fig. 4C). Notably, there were no significant differences in the levels of TNFα, IL6, IL10 or TGFβ in the testes of LPS/PG/polyIC-treated mice compared to saline controls, 8 weeks after exposure (Figs. 4D, 4F-H). However, there was a marginal increase in IL1β levels in PG and polyIC treated mice (Fig. 4E).

**Figure 4.**
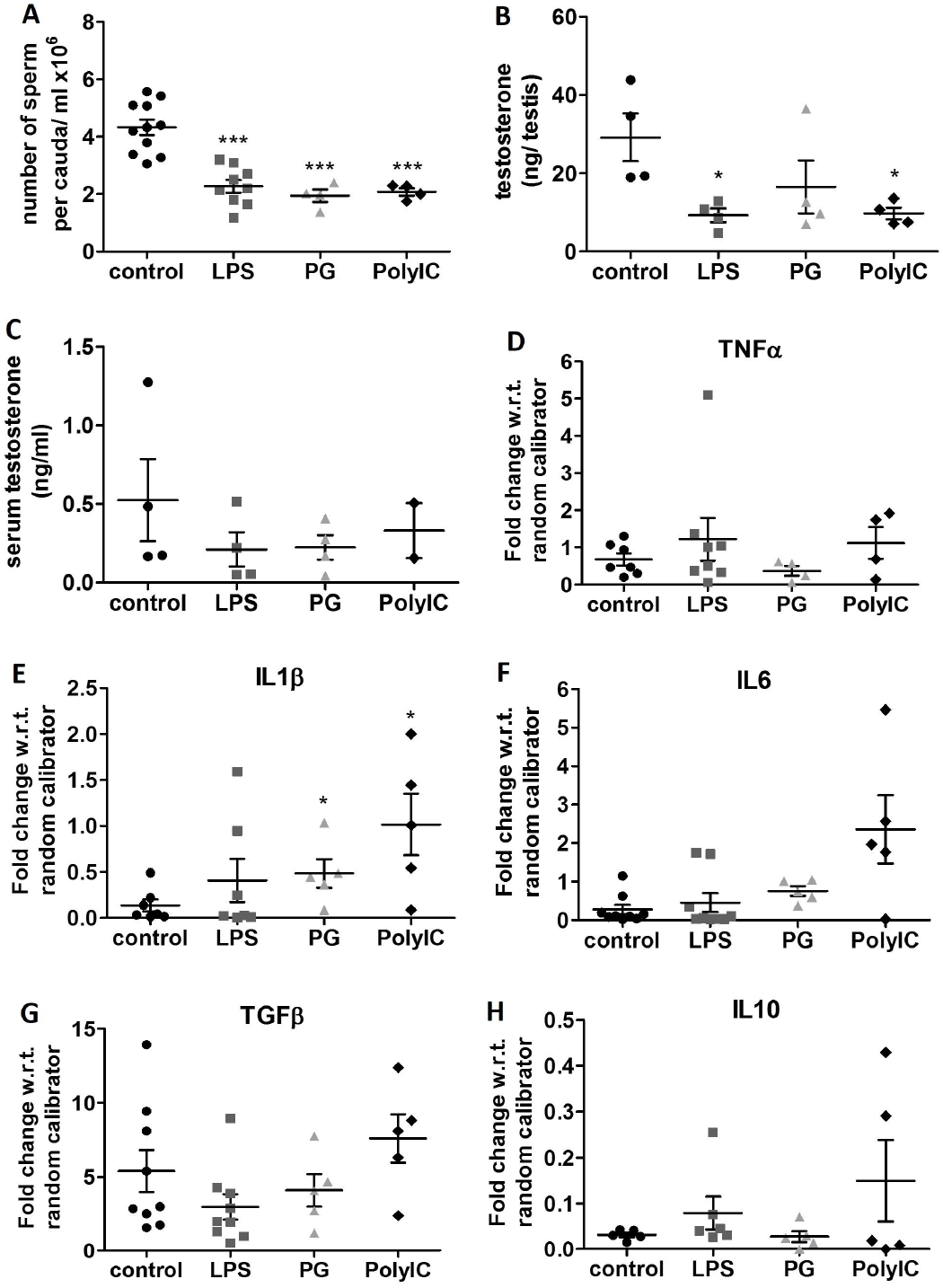
Long term impact of a single transient systemic TLR agonist exposure on sperm counts, testosterone levels and testicular cytokine milieu in mice. Eight week after a single i.p. injection of LPS/PG/polyIC, (A) caudal sperm counts (n = 4-11), (B) testicular testosterone levels (n = 4) and (C) serum testosterone levels (n = 2-were measured. (D-H) Levels of mRNA for TNFα (D), IL1β (E), IL6 (F), TGFβ (G), IL10 (H) were measured using qRTPCR (n = 4-8). Data represent mean ± s.e.. *p< 0.05, ***p<0.0001.

### Effects of a single LPS/PG/polyIC exposure on progeny sired 8 weeks later

We next tested if these features of testicular dysfunction affected the fertility potential of LPS/PG/polyIC-exposed male mice. For this, 8 weeks after LPS, PG, polyIC or saline injection, each male was individually cohabited for 3 weeks with sexually mature healthy female mice of proven fertility. Pregnancies resulted from all matings set up, with either saline-treated or LPS/PG/polyIC-treated sires. The resultant litter sizes were also comparable between the various groups of sires (Fig. 5A). However, the neonates from LPS-treated sires showed substantially lower birth weights than those from saline-treated sires (Fig. 5B). Notably, this effect was seen with PG-treated and polyIC-treated sires as well (Fig. 5B).

**Figure 5.**
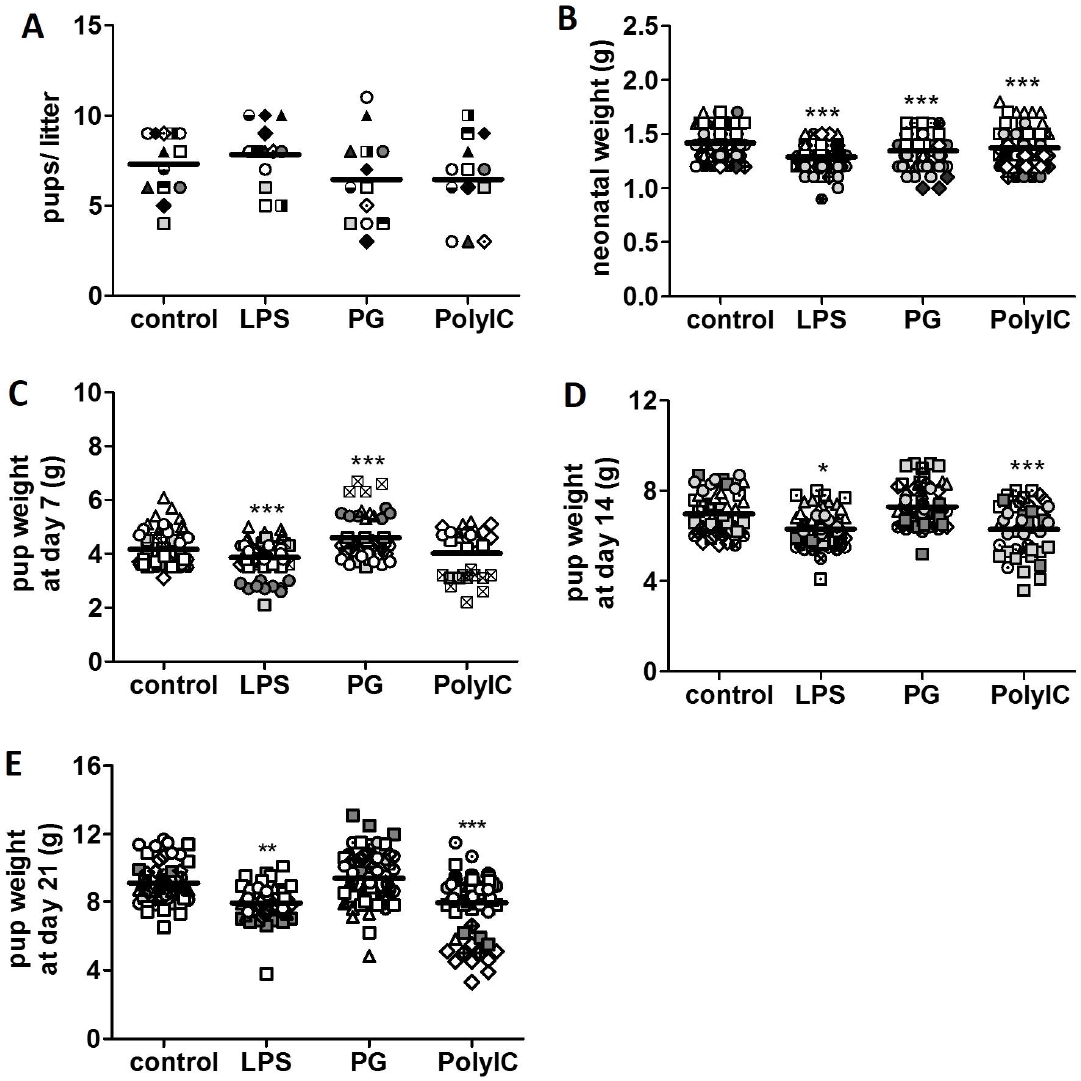
Low neonatal body weights and poor weight gain in progeny sired by mice treated with LPS/PG/polyIC eight weeks previously. Data on progeny from proven fertile females mated with male mice treated 8 weeks earlier with LPS, PG or polyIC or saline (control). (A) Number of pups in each litter (n = 13 litters). (B) Pup weights at birth (n = 13-15 litters representing 84-116 pups). (C) Pup weights at post-natal day 7 (n = 6-9 litters representing 29-62 pups). (D) Pup weights at postnatal day 14 (n = 8-11 litters representing 45-70 pups). (E) Pup weights at day 21 (n = 6-12 litters representing 48-70 pups). Each litter is represented by a distinct symbol. *p< 0.05, **p<0.01, ***p<0.0001.

There is evidence that low-birth weight neonates can undergo compensatory weight gain, particularly in adipose tissue, and this phenotype is accompanied by an insulin-resistant hyperglycemic metabolic syndrome-like phenotype (24). We therefore followed the weight gain of these neonates over the next three weeks. It was apparent that progeny from LPS-treated sires remained smaller all through this period (Figs. 5C-5E). Progeny from polyIC-treated sires tended to remain small over this period, but progeny from PG-treated sires showed comparatively higher average weight than the control group (Figs. 5C-5E). At 8 weeks of age, male progeny mice showed no differences in their lean weights or their lengths (Figs. 6A-6B). However, female progeny mice showed marginally lower lean body weight (Fig. 6D) but not in length (Fig. 6E). When serum glucose levels were estimated in them, they showed no differences between the various groups of progeny (Figs. 6C, 6F).

**Figure 6.**
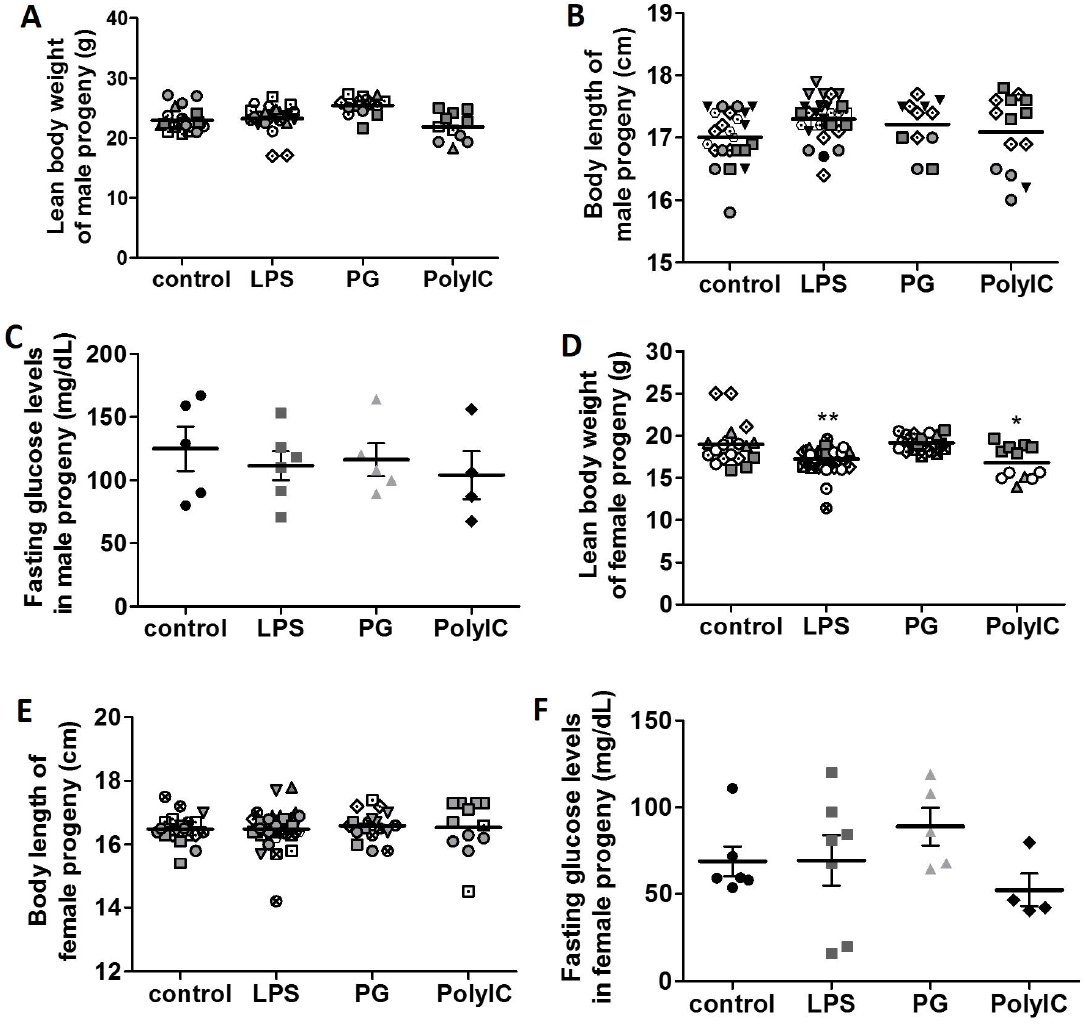
Absence of metabolic differences in adult progeny sired by mice treated with LPS/PG/polyIC eight weeks previously. Data on 8 week-old progeny from proven fertile females mated with male mice treated 8 weeks earlier with LPS, PG or polyIC or saline (control). (A) Lean body weights of male progeny. (B) Body length of male progeny. (C) Fasting blood glucose levels of male progeny after 16 h of fasting. (D) Lean body weights of female progeny. (E) Body length of female progeny. (F) Fasting blood glucose levels of female progeny after 16 h of fasting. For panels A, B, D and E, each litter is represented by a different symbol. For glucose estimation n = 4-7. For other panels n ≥ 12. Mean ± s.e. is shown. *p< 0.05, **p<0.01.

In order to examine the possible basis of the lower birth weight further, pregnant females were euthanized at 17.5 dpc and fetal lengths and fetal and placental weights were recorded. Placental weights were clearly lower in the groups with LPS/PG/polyIC-treated sires (Fig.7A) whereas fetal weights and lengths showed no differences between progeny of LPS/PG/polyIC-treated versus saline-treated sires (Figs. 7B-7C).

**Figure 7.**
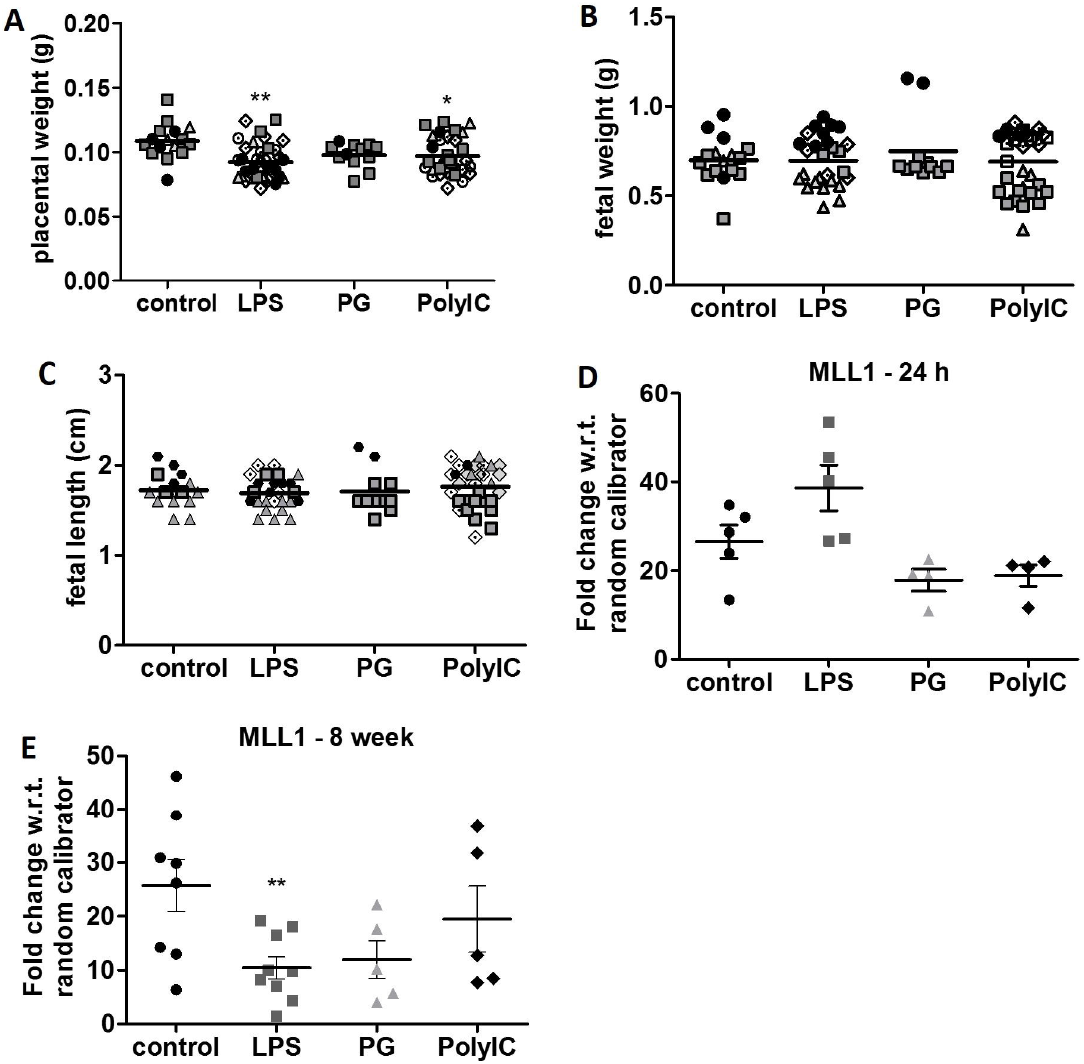
Low placental weights in pregnancies sired by mice treated with LPS/PG/polyIC eight weeks previously, with evidence of testicular epigenetic modulation in these sires. Proven fertile females were mated with sires treated 8 weeks earlier with LPS, PG or polyIC, or saline (control). Occurrence of vaginal plug was treated as day 0.5 of pregnancy. At 17.5 days, pregnant dams were euthanized to measure (A) placental weights, (B) fetal weights, and (C) fetal lengths (n = 12-33 placentae or fetuses from 2-5 dams). Data from each dam are represented by a different symbol. (D, E) Normalised transcript levels of MLL1 in the testes, 24 h (D, n = 4-5) and 8 weeks (E, n = 5-9) after treatment with LPS, PG, polyIC or saline (control). Data represent mean ± s.e.. *p< 0.05, **p<0.01.

These data suggested that the epigenetic landscape of sperm from LPS/PG/polyIC-treated mice was likely to have been altered with reference to its post-fertilization programming for placental formation. We therefore tested the testicular levels of a major modifier of epigenetic programming that is known to respond to inflammatory stimuli, the H3K4-specific histone methyl-transferase enzyme mixed-lineage leukemia-1 (MLL1). While MLL1 levels in the testes of LPS/PG/polyIC-treated mice did not show changes at 24 h after administration (Fig. 7D), by 8 weeks after exposure, testicular MLL1 levels were substantially reduced (Fig. 7E).

## Discussion

Our data here show that there are long-term trans-generational functional consequences of even a single transient systemic inflammatory event in the testes. While the resultant local testicular inflammation is transient and subsides within a week, it leads to long-term alteration of the spermatogenetic programme. This results in a modest reduction of sperm counts and epigenetic alterations in post-fertilisation events such as placentogenesis and progeny weight.

Immune privilege in tissues is thought to be mediated by a blood-tissue barrier formed by local endothelial cell properties, ensuring that diverse blood-borne activators, cells as well as molecules, do not easily enter the privileged tissues (25, 26). One consequence of such sequestration is prevention of access of potentially auto-reactive T cells to tissue-specific sequestered self-antigenic proteins (27, 28). Another consequence is the prevention of easy ingress of infectious agents as well as of inflammatory cells (29). Nonetheless, there are examples of tissue-tropic infections for the testes, such as infections by *Chlamydia trachomatis* or uropathogenic *Escherichia coli*; or by viruses like mumps virus, Coxsackie virus, or zika virus (30, 31, 32). Similarly, there is evidence that the testes have the capacity to mediate local inflammatory responses. Testicular cells express an array of PRRs such as TLRs, RIG-I-like receptors and NOD-like receptors, which are actively involved in mediating testicular innate immune responses (14, 15, 16, 17, 33). Further, the persistence of a number of viruses in the testes and epididymides of male mice has been shown to be associated with induction of pro-inflammatory mediators, tissue damage, diminished testosterone levels, impaired blood–testis barrier (BTB) and sperm quality (7, 8, 9, 10, 34).

Our findings at early time points upon systemic TLR ligand administration were consistent with earlier reports (18, 20, 35), by showing local induction in the testes of mRNA for a number of cytokines; TNFα, IL1β and IL6. Notably, TLR2 and TLR3 ligands were less potent than TLR4 activation in inducing local inflammatory responses in the testes. The next issue to examine was the extent, if any, to which this transient local inflammation modulated testicular function. Activation of TLR3 or TLR4 has been shown to suppress steroidogenesis by Leydig cells (16). Also, sperm vitality was reduced at early time points in rats given a single dose of systemic LPS (36). Consistent with these findings, our data showed that caudal sperm counts were significantly decreased within 24 h of systemic LPS injection, though not with PG or polyIC. However, both the local cytokine responses and the sperm count reduction came back to normal baselines within a week of induction of transient systemic inflammation. This was not only true of the testes but of the brain as well. There were local inflammatory responses that returned to baseline within a few days, and the acute functional consequences were seen only on the first day after TLR ligand administration.

Under these circumstances, there appeared no prior reason to expect any long-term testicular consequences of this transient inflammation. However, there is evidence that, five weeks after a single systemic dose of LPS, male mice can show continuing features of persistent inflammation with infiltration of immune cells in the interstitial spaces, thickening of smooth muscle layers, and dysregulated spermatogenesis with loss of germ cells in seminiferous tubules and presence of germ cells in the caudal lumen (22). Our data did not show any persistent local testicular inflammation at 8 weeks after LPS/PG/polyIC-exposure. However, we did observe a modest but consistent reduction in the caudal sperm count 8 weeks after LPS/PG/polyIC-exposure, accompanied by a similarly modest but consistent reduction in testosterone levels.

Thus, our data suggest a trajectory in which there is early transient local inflammation in the testes accompanied by reduction of sperm counts, which resolves within a week, and yet leads to a reduction in sperm counts and testosterone levels 8 weeks later without any persistent inflammation. While TLR2 or TLR3 activation did not lead to as intense inflammation or sperm count reduction as TLR4 activation did, the late alterations in testicular function were almost equivalent in all three cases. Therefore, it is possible either that the late events were not in fact mediated by the early inflammatory events but by an independent TLR-mediated pathway, and/or that the late dysfunctions were far more sensitive than the early events were. The fact that both Leydig cells and germ cells express TLRs (15, 16) may be relevant in this context.

Persistent reduction of caudal sperm counts could plausibly be related to germ cell damage, including some apoptotic loss. Since the range of apparently normal sperm counts is wide, a two-fold difference would seem unlikely to result in any detectable loss of fecundity, and in fact, mating studies did not reveal any deficits in the fecundity of TLR agonist-treated male mice. Studies with male mice carrying targeted deletions of a number of genes with important roles in testicular spermatogenesis and steroidogenesis have been reported to exhibit normal fertility even with significantly reduced sperm counts or testicular weights (37, 38, 39). The female genital tract is known to exert strong selection pressure to promote movement of a small but high-quality fertilising subpopulation of spermatozoa to ultimately reach the fertilisation site (40). Therefore, deficits in sperm counts or function could be masked by the proficiency of this selection system.

However, since the TLR agonist-treated male mice showed evidence of an altered programme of spermatogenesis, we further examined the pregnancies for any evidence of dysregulated embryogenesis. Clearly, transient paternal exposure to TLR agonists led to defects in placental and fetal growth *in utero*. The placenta plays a key role in fetal nutrition, via active transport of nutrients and metabolic byproducts across the feto-maternal interface (41), and changes in placental nutrient transport can directly contribute to altered fetal growth (42). Our data suggest that transient exposure to TLR ligands led to long-term epigenetic alterations in the sperm that caused poor placental maturation.

It has become increasingly clear that sperm can transfer epigenetic information to influence fetal phenotypes (43), and a number of examples of such modifications have been reported in both animal models and in the human system (44, 45, 46, 47). Paternal dietary modifications have been extensively reported to alter progeny metabolic phenotypes (46, 48, 49). Similarly, paternal stress has been shown to affect a number of progeny phenotypes (50, 51). There is also growing evidence that paternal epigenetic influences modify the development and maturation of the placenta and can thus lead to underweight progeny (52, 53).

Epigenetic alterations are thus potential determinants that link paternal environment to sperm quality and offspring development. The mechanisms of epigenetic alterations in sperm that affect fetal outcomes are thus of interest. The molecular pathways shown to be able to mediate epigenetic signatures in sperm include sperm RNAs, DNA methylation as well as histone modifications (46, 48, 54). Studies with male mice fed high–fat or low-protein diets have demonstrated an increase in the levels of fragmented small tRNAs (stRNA) in sperm and altered patterns of sperm DNA methylation, associated with metabolic diseases in offspring (48, 55). When male mice maintained on restricted diet for a prolonged period of 10 weeks were mated to untreated females, they sired offspring with reduced postnatal weight and growth but increased adiposity and dyslipidaemia (56). Importantly, another study of transient starvation model where animals were subjected to transient 24 h fasting, when mated to untreated females 2 weeks later, sired offspring exhibiting decreased serum glucose levels. However, transient fasting of sires in this study, did not significantly alter the average litter size, sex ratio and body weight of the offspring (57). Findings from these reports imply that, minor body weight reduction observed transiently upon LPS or polyIC injection, may not be the reason for low birth weight progenies observed in our study. This notion is further supported by the fact that, the PG challenged sires in our study though did not exhibit any body weight reduction, sired progenies with low birth weight. Immune activation via polymicrobial abdominal sepsis in sires was shown to affect progeny immune phenotypes, accompanied by differentially hypo- and hyper-methylated cytosine residues spread over the sperm genome, implicating the sperm methylome as a potential carrier of epigenetic information (58). Further, paternal activation of CB2 cannabinoid receptor led to impaired placental and embryonic growth via altered DNA methylation/hydroxymethylation levels at imprinted genes in sperm (53).

Our data so far provide no direct evidence of the precise epigenetic pathways altered in sperm developing eight weeks after a single transient systemic inflammatory episode. We examined the testicular expression of a potential marker that would indicate epigenetic alteration in the testes subsequent to inflammation. Mixed-lineage leukemia-1 (MLL1), a histone methyl-transferase enzyme with site specificity for H3K4, is induced by inflammatory signals in a number of cell lineages (59, 60). It is known to be required for the epigenetic maintenance of gene activation during development (61, 62), and disruption of H3K4 methylation in sperm has been reported to have transgenerational consequences (54). Indeed, our findings of decreased levels of MLL1 in the testes of TLR agonist-injected mice 8 weeks after exposure show that major epigenetic modification of sperm programming is likely to have taken place. The precise pathways involved thus become future directions of major interest.

Underweight human neonates that gain weight postnatally in compensatory fashion have been shown to acquire a more adipose phenotype, and to be more prone to early acquisition of insulin resistance (63, 64). We therefore followed the postnatal weights of the underweight progeny mice almost to the end of the weaning period and tested their lean weights, lengths, and blood glucose levels at early adulthood. The underweight progeny of LPS- or polyIC-treated sires remained underweight, while those of PG-treated sires tended to recover weight. However, during later periods, the lean weights of progenies were similar, suggesting that there was no increase in adiposity, and blood glucose levels were comparable, indicating that, in this model of intra-uterine growth retardation, postnatal metabolic syndrome was not a prominent feature.

In conclusion, our findings demonstrate that exposure to transient systemic inflammation has persistent sperm-mediated consequences on progeny health. Maternal health, especially in low-income communities with high levels of exposure to infectious diseases, is widely linked to neonatal health, especially to intra-uterine growth retardation (65). However, attempts to improve these neonatal weight outcomes by improved maternal health and nutrition are only partly successful (66, 67, 68). Our data indicate that paternal high-frequency pathogen exposure may well be a factor contributing epigenetically to the developing country problem of frequent intra-uterine growth retardation.

## Acknowledgements

Financial support from the DBT-RA Program in Biotechnology and Life Sciences Department of Biotechnology, Government of India to SuR and from Cognitive Science Research Initiative, Department of Science and Technology, Government of India to AG and NS (Grant No. SR/CSRI/331/2016(G)) is gratefully acknowledged. Indian Institute of Science Education and Research, Pune is funded by Ministry of Human Resource Development, Government of India.

## Conflict of interest

The authors declare no conflict of interest.

## Notes

### Competing Interest Statement

The authors have declared no competing interest.

## References

1. Ravindranath N, Dettin L, and Dym M. Mammalian Testes: Structure and Function. Springer, 2003;1–19.

2. Griswold MD. Spermatogenesis: The current commitment to meiosis. Physiol Rev. 2016; 96(1):1–17.

3. Sharma U. Paternal contributions to offspring health: Role of sperm small RNAs in intergenerational transmission of epigenetic information. Front Cell Dev Biol. 2019;7:215.

4. Mruk DD, Cheng CY. The Mammalian Blood-Testis Barrier: Its Biology and Regulation. Endocrine Reviews. 2015;36(5):564–591.

5. Meinhardt A, Hedger MP. Immunological, paracrine and endocrine aspects of testicular immune privilege. Mol Cell Endocrinol. 2011;335(1):60–68.

6. Pellati D, Mylonakis I, Bertoloni G, Fiore C, Andrisani A, Ambrosini G, Armanini D. Genital tract infections and infertility. Eur J Obstet Gynecol Reprod Biol. 2008;140(1):3–11.

7. Shan C, Muruato AE, Jagger BW, et al. A single–dose live-attenuated vaccine prevents Zika virus pregnancy transmission and testis damage. Nat Commun. 2017;8(1):676

8. Matusali G, Dereuddre-Bosquet N, Le Tortorec A, Moreau M, Satie A-P, Mahé D, Roumaud P, Bourry O, Sylla N, Bernard-Stoecklin S, Pruvost A, Le Grand R, Dejucq-Rainsford N. Detection of simian immunodeficiency virus in semen, urethra, and male reproductive organs during efficient highly active antiretroviral therapy. J Virol. 2015;89:5772–5787.

9. Malolina EA, Kulibin AY, Naumenko VA, Gushchina EA, Zavalishina LE, Kushch AA. Herpes simplex virus inoculation in murine rete testis results in irreversible testicular damage. Int J Exp Path. 2014;95(2):120–130.

10. Le Tortorec A, Le Grand R, Denis H, Satie A-P, Mannioui K, Roques P, Maillard A, Daniels S, Jégou B, Dejucq-Rainsford N. Infection of semen-producing organs by SIV during the acute and chronic stages of the disease. PLoS One. 2008;3(3): e1792.

11. Zhao S, Zhu W, Xue S, Han D. Testicular defense systems: immune privilege and innate immunity. Cell Mol Immunol. 2014;11(5): 428–437.

12. Schuppe HC, Meinhardt A, Allam JP, Bergmann M, Weidner W, Haidl G. Chronic orchitis: a neglected cause of male infertility? Andrologia. 2008;40(2):84–91.

13. Li N, Wang T, Han D. Structural, cellular and molecular aspects of immune privilege in the testis. Front Immunol. 2012;3:152.

14. Zhu W, Chen Q, Yan K, Liu Z, Li N, Zhang X, Yu L, Chen Y, Han D. RIG-I-like receptors mediate innate antiviral response in mouse testis. Mol Endocrinol. 2013; 27(9):1455–1467.

15. Wang T, Zhang X, Chen Q, Deng T, Zhang Y, Li N, Shang T, Chen Y, Han D. Toll-like receptor 3-initiated antiviral responses in mouse male germ cells in vitro. Biol Reprod. 2012;86(4):106.

16. Shang T, Zhang X, Wang T, Sun B, Deng T, Han D. Toll-like receptor-initiated testicular innate immune responses in mouse Leydig cells. Endocrinology. 2011; 152(7):2827–2836.

17. Winnall WR, Muir JA, Hedger MP. Differential responses of epithelial Sertoli cells of the rat testis to Toll-like receptor 2 and 4 ligands: implications for studies of testicular inflammation using bacterial lipopolysaccharides. Innate Immun 2011;17:123–136.

18. Rokade S, Kishore U, Madan T. Surfactant Protein D regulates murine testicular immune milieu and sperm functions. Am J Reprod Immunol. 2017;77(3).

19. Sarkar O, Bahainwala J, Chnadrasekaran S, Kothari S, Mathur PP, Agarwal A. Impact of inflammation on male fertility. Front Biosci (Elite Ed). 2011;3:89–95.

20. O’Bryan MK, Gerdprasert O, Nikolic-Paterson DJ, Meinhardt A, Muir JA, Foulds LM, Phillips DJ, de Kretser DM, Hedger MP. Cytokine profiles in the testes of rats treated with lipopolysaccharide reveal localized suppression of inflammatory responses. Am J Physiol Regul Integr Comp Physiol. 2005;288(6):R1744–55.

21. O’bryan MK, Schlatt S, Phillips DJ, De Kretser DM, Hedger MP. Bacterial lipopolysaccharide-induced inflammation compromises testicular function at multiple levels *in vivo*. Endocrinology. 2000;141:238–246.

22. Jafari O, Babaei H, Kheirandish R, Samimi A-S, Zahmatkesh A. Histomorphometric evaluation of mice testicular tissue following short- and long-term effects of lipopolysaccharide-induced endotoxemia. Iran J Basic Med Sci. 2018;21(1):47–52.

23. Wang F, Liu W, Jiang Q, Gong M, Chen R, Wu H, Han R, Chen Y, Han D. Lipopolysaccharide-induced testicular dysfunction and epididymitis in mice: a critical role of tumor necrosis factor alpha†. Biol Reprod. 2019;100(3):849–861.

24. Symonds M E, Pope M, Sharkey D, Budge H. Adipose tissue and fetal programming. Diabetologia. 2012;55:1597–1606.

25. Mital P, Hinton BT, Dufour JM. The blood-testis and blood-epididymis barriers are more than just their tight junctions. Biol Reprod. 2011;84(5):851–858.

26. Bechmann I, Galea I, Perry V H. What is blood-brain barrier (not)? Trends Immunol. 2007;28(1):5–11.

27. Dym M, Romrell LJ. Intraepithelial lymphocytes in the male reproductive tract of rats and rhesus monkey. J Reprod Fertil. 1975;42:1–7.

28. Setchell BP, Voglmayr JK, Waites GMH. A blood-testis barrier restricting passage from blood into rete testis fluid but not into lymph. J Physiol. 1969;200(1):73–85.

29. Cheng CY, Mruk DD. The blood-testis barrier and its implications for male contraception. Pharmacol Rev. 2012;64(1):16–64.

30. Wu H, Jiang X, Gao Y, Liu W, Wang F, Gong M, Chen R, Yu X, Zhang W, Gao B, Song C, Han D. Mumps virus infection disrupts blood-testis barrier through the induction of TNF-α in Sertoli cells. FASEB J. 2019;33(11):12528–12540.

31. Fijak M, Pilatz A, Hedger MP, Nicolas N, Bhushan S, Michel V, Tung KSK, Schuppe HC, Meinhardt A. Infectious, inflammatory and ‘autoimmune’ male factor infertility: how do rodent models inform clinical practice? Hum Reprod Update. 2018;24(4):416–441.

32. Siemann DN, Strange DP, Maharaj PN, Shi PY, Verma S. Zika virus infects human Sertoli cells and modulates the integrity of the *in vitro* blood-testis barrier model. J Virol. 2017;91(22):e00623–17.

33. Starace D, Galli R, Paone A, De Cesaris P, Filippini A, Ziparo E, Riccioli A. Toll-like receptor 3 activation induces antiviral immune responses in mouse Sertoli cells. Biol Reprod. 2008; 79:766–775.

34. Wu H, Shi L, Wang Q, Cheng L, Zhao X, Chen Q, Jiang Q, Feng M, Li Q, Han D. Mumps virus-induced innate immune responses in mouse Sertoli and Leydig cells. Sci Rep 6, 19507 (2016).

35. Palladino MA, Fasano GA, Patel D, Dugan C, London M. Effects of lipopolysaccharide-induced inflammation on hypoxia and inflammatory gene expression pathways of the rat testis. Basic Clin Androl. 28, 14(2018).

36. Hassan A, Youssef M, Khalil AM, Ahmed H. Effects of bacterial lipopolysaccharide on serum testosterone level and sperm vitality in mature rats. Journal of Veterinary Medical Research. 2017;24(2):163–168.

37. Enright BP, Davila DR, Tornesi BM, Blaich G, Hoberman AM, Gallenberg LA. Developmental and reproductive toxicology studies in IL-12p40 knockout mice. Birth Defects Res B Dev Reprod Toxicol. 2011;92(2):102–110.

38. Schürmann A, Koling S, Jacobs S, Saftig P, Krauss S, Wennemuth G, Kluge R, Joost HG. Reduced sperm count and normal fertility in male mice with targeted disruption of the ADP-ribosylation factor-like 4 (Arl4) gene. Mol Cell Biol. 2002;22(8):2761–2768.

39. Kumar TR, Wang Y, Lu N, Matzuk MM. Follicle stimulating hormone is required for ovarian follicle maturation but not male fertility. Nat Genet. 1997;15(2):201–204.

40. Sakkas D, Ramalingam M, Garrido N, Barratt CL. Sperm selection in natural conception: what can we learn from Mother Nature to improve assisted reproduction outcomes? Hum Reprod Update. 2015;21(6):711–26.

41. Garnica AD, Chan WY. The role of the placenta in fetal nutrition and growth. J Am Coll Nutr. 1996;15(3):206–222.

42. Lager S, Powell TL. Regulation of nutrient transport across the placenta. Journal of Pregnancy. 2012. Article ID 179827.

43. Rando OJ. Intergenerational transfer of epigenetic information in sperm. Cold Spring Harb Perspect Med. 2016;6(5):a022988.

44. Soubry A, Murphy S K, Wang F, Huang Z, Vidal AC, Fuemmeler BF, Kurtzberg J, Murtha A, Jirtle RL, Schildkraut JM, Hoyo C. Newborns of obese parents have altered DNA methylation patterns at imprinted genes. Int J Obes. 2015;39(4):650–657.

45. Lambrot R, Xu S, Saint-Phar S, Chountalos G, Cohen T, Paquet M, Suderman M, Hallett M, Kimmins S. Low paternal dietary folate alters the mouse sperm epigenome and is associated with negative pregnancy outcomes. Nat Commun. 2013;4:4:2889.

46. Fullston T, Ohlsson Teague EMC, Palmer N O, DeBlasio M J, Mitchell M, Corbett M, Print C G, Owens J A, Lane M. Paternal obesity initiates metabolic disturbances in two generations of mice with incomplete penetrance to the F2 generation and alters the transcriptional profile of testis and sperm microRNA content. FASEBJ. 2013;27:4226–4243.

47. Meek LR, Myren K, Sturm J, Burau D. Acute paternal alcohol use affects offspring development and adult behaviour. Physiol Behav. 2007;91(1):154–60.

48. Carone BR, Fauquier L, Habib N, Shea JM, Hart CE, Li R, Bock C, Li C et al. Paternally induced transgenerational enviornmental reprogramming of metabolic gene expression in mammals. Cell. 2010;143(7):1084–1096.

49. Ng S-F, Lin RC, Laybutt DR, Barres R, Owens JA, Morris MJ. Chronic high-fat diet in fathers programs β-cell dysfunction in female rat offspring. Nature. 2010;467(7318):963–966.

50. Gapp K, Jawaid A, Sarkies P, Bohacek J, Pelczar P, Prados J, Farinelli L, Miska E, Mansuy IM. Implication of sperm RNAs in transgenerational inheritance of the effects of early trauma in mice. Nat Neurosci. 2014;17(5):667–669.

51. Rodgers AB, Morgan CP, Bronson SL, Revello S, Bale TL. Paternal stress exposure alters sperm microRNA content and reprograms offspring HPA stress axis regulation. J Neurosci. 2013;33(21):9003–9012.

52. Denomme M M, Parks J C, McCallie B R, McCubbin N I, Schoolcraft W B, Katz-Jaffe M G. Advanced paternal age directly impacts mouse embryonic placental imprinting. PLoS ONE. 2020;15(3): e0229904.

53. Innocenzi E, Domenico ED, Ciccarone F, Zampieri M, Rossi G, Cicconi R, Bernardini R, Mattei M, Grimaldi P. Paternal activation of CB_2_ cannabinoid receptor impairs placental and embryonic growth via epigenetic mechanism. Sci Rep 9, 17034 (2019).

54. Siklenka K, Erkek S, Godmann M, Lambrot R, McGraw S, Lafleur C, Cohen T, Xia J, Suderman M, Hallett M, Trasler J, Peters AH, Kimmins S. Disruption of histone methylation in developing sperm impairs offspring health transgenerationally. Science. 2015;350(6261):aab2006.

55. Chen Q, Yan W, Duan E. Epigenetic inheritance of acquired traits through sperm RNAs and sperm RNA modifications. Nat Rev Genet. 2016;17(12):733–743.

56. McPherson N, Fullston, T, Kang W, Sandeman L Y, Corbett M A, Owens J A, Lane M. Paternal under-nutrition programs metabolic syndrome in offspring which can be reversed by antioxidant/vitamin food fortification in fathers. Sci Rep 6,27010 (2016).

57. Anderson L M, Riffle L, Wilson R, Travlos G S, Lubomirski M S, Alvord W G. Preconceptional fasting of fathers alters serum glucose in offspring of mice. Nutrition. 2006;22(3):327–331.

58. Bomans K, Schenz J, Tamulyte S, Schaack D, Weigand MA, Uhle F. Paternal sepsis induces alterations of the sperm methylome and dampens offspring immune responses-an animal study. Clin Epigenetics. 2018;10:89.

59. Kimball A S, Joshi A, Carson WF 4th, Boniakowski AE, Schaller M, Allen R, Bermick J, Davis FM, Henke PK, Burant CF, Kunkel SL, Gallagher KA. The histone methyltransferase MLL1 directs macrophage-mediated inflammation in wound healing and is altered in a murine model of obesity and type 2 diabetes. Diabetes. 2017;66(9):2459–2471.

60. Carson WF 4th, Cavassani KA, Soares EM, Hirai S, Kittan NA, Schaller M A, Scola MM, Joshi A et al. The STAT4/MLL1 epigenetic axis regulates the antimicrobial functions of murine macrophages. J Immunol. 2017;199(5):1865–1874.

61. Lim DA, Huang Y-C, Swigut T, Mirick AL, Garcia-Verdugo JM, Wysocka J, Ernst P, Alvarez-Buylla A. Chromatin remodelling factor MLL1 is essential for neurogenesis from postnatal neural stem cells. Nature. 2009;458(7237):529–533.

62. Jude CD, Climer L, Xu D, Artinger E, Fisher JK, Ernst P. Unique and independent roles for MLL in adult hematopoietic stem cells and progenitors. Cell Stem Cell. 2007;1(3):324–337.

63. Ferrannini E, Iozzo P, Virtanen KA, Honka M-J, Bucci M, Nuutila P. Adipose tissue and skeletal muscle insulin-mediated glucose uptake in insulin resistance: Role of blood flow and diabetes. Am J Clin Nutr. 2018;108(4):749–758.

64. Yajnik CS. Nutrient-mediated teratogenesis and fuel-mediated teratogenesis: Two pathways of intrauterine programming of diabetes. Int Gynaecol Obstet. 2009;104 Suppl 1:S27–31.

65. Malhotra A, Allison B J, Castillo-Melendez M, Jenkin G, Polglase G R, Miller S L. Neonatal morbidities of fetal growth restriction: pathophysiology and impact. Front Endocrinol. 2019;10:55.

66. Hambidge KM, Krebs NF. Strategies for optimizing maternal nutrition to promote infant development. Reprod Health 15, 87 (2018).

67. Mori R, Ota E, Middleton P, Tobe-Gai R, Mahomed K, Bhutta ZA. Zinc supplementation for improving pregnancy and infant outcome. Cochrane Database Syst Rev. 2012;7:CD000230.7

68. Imdad A, Bhutta ZA. Maternal nutrition and birth outcomes: effect of balanced protein-energy supplementation. Paediatric and Perinatal Epidemiology. 2012;26 (Suppl. 1):178–90.

