## Supplementary figure legends for "Transient systemic inflammation in adult male mice results in underweight progeny"

***Figure S1: Open field arena behaviour analysis after saline (control) or LPS administration***

Representative trails from control and LPS-treated groups of mice at different time intervals.

***Figure S2: Exploratory behaviour of mice challenged with LPS in open field test***

Compiled data from saline or LPS injected mice subjected to open field test analysed using EthoVision software. (A) Mean velocity of animal locomotion in cm/s, (B) total distance moved by the animal in the area in cm, (C) total number of rearings measured manually, (D) total time spent in the periphery of the area in sec, (E) total time spent in the centre of the area in sec. (n= 3 mice per group). Data represent mean ± s.e.. *p < 0.05.

***Figure S3: Cytokine responses in the spleens of LPS/ PG/polyIC injected mice***

Normalised mRNA levels of TNFα (A), IL1β (B), IL6 (C), TGFβ (D) and IL10 (E) using qRTPCR from spleens of mice 24 h after LPS/PG/polyIC or saline administration (n =2-5 mice per group). Data shown as mean ± s.e..

***Figure S4: Cytokine responses in the brains of LPS treated mice***

Normalised mRNA levels of TNFα (A), IL1β (B), IL6 (C), IL10 (D), and TGFβ (E) using qRTPCR at indicated times from brains of mice after treatment with LPS or saline (n- 3-5 mice per group). Data are shown as mean ± s.e.. *p< 0.05, **p<0.01.

***Figure S5:*** ***Open field arena behaviour analysis after saline (control) or PG administration***

Representative trails from control and PG-treated groups of mice at different time intervals.

***Figure S6:*** ***Exploratory behaviour of mice challenged with PG in open field test***

Compiled data from saline or PG-treated mice subjected to open field test analysed using EthoVision software. (A) Mean velocity of animal locomotion in cm/s, (B) total distance moved by the animal in the area in cm, (C) total number of rearings measured manually, (D) total time spent in the periphery of the area in sec, (E) total time spent in the centre of the area in sec. (n = 3). Data shown as mean ± s.e..

***Figure S7: Open field arena behaviour analysis after saline (control) or polyIC administration***

Representative trails from control and polyIC injected groups of mice at different time intervals.

***Figure S8:*** ***Exploratory behaviour of mice challenged with polyIC in open field test***

Compiled data from saline or polyIC-treated mice subjected to open field test analysed using EthoVision software. A) Mean velocity of animal locomotion in cm/s, B) total distance moved by the animal in the area in cm, C) total number of rearings measured manually, D) total time spent in the periphery of the area in sec, E) total time spent in the centre of the area in sec. (n=3). Data shown as mean ± s.e..

***Figure S9:*** ***Cytokine responses in the brains of PG-treated mice***

After PG injection i.p., mRNA levels of TNFα (A), IL1β (B), IL6 (C), IL10 (D) and TGFβ (E) were measured in brains by qRTPCR and normalised values are shown at indicated times (n= 3-5). Data represent mean ± s.e..

***Figure S10:*** ***Cytokine responses in the brains of polyIC-treated mice***

After polyIC injection i.p., mRNA levels of TNFα (A), IL1β (B), IL6 (C), IFNγ (D), IL10 (E) and TGFβ (F) were measured in brains by qRTPCR and normalised values are shown at indicated times. (n=3-5). Data represent mean ± s.e..
