## Supplementary table for "Transient systemic inflammation in adult male mice results in underweight progeny"

***Table S1: Details of mouse primer sequences used***

|  | **Target gene** | **Annealing**  **temperature** | **Product size**  **(base pairs)** | **5’→3’ sequences**  **(F: forward primer;**  **R: reverse primer)** |
| --- | --- | --- | --- | --- |
| 1 | TNFα^ǂ^ | 64ºC | 237 | F: GGGCCACCACGCTCTTCTGTCT  R: GATCCATGCCGTTGGCCAGGAG |
| 2 | TGFβ^ǂ^ | 68ºC | 130 | F: GCGTGCTAATGGTGGACCGCA  R: GCACGGGACAGCAATGGGGG |
| 3 | IL1β^ǂ^ | 70ºC | 139 | F: GCCTCGTGCTGTCGGACCCAT  R: TTGAGGCCCAAGGCCACAGGTA |
| 4 | IL10^ǂ^ | 68ºC | 158 | F: GCACCCACTTCCCAGTCGGC  R: GGCTTGGCAACCCAAGTAACCCT |
| 5 | IL6^ǂ^ | 64ºC | 146 | F: AGACAAAGCCAGAGTCCTTCAGAGA  R: GCCACTCCTTCTGTGACTCCAGC |
| 6 | IFNγ | 64ºC | 162 | F: TTCTTCAGCAACAGCAAGGCGA  R: GCAGCGACTCCTTTTCCGCT |
| 7 | MLL1 | 60ºC | 271 | F: CTTCTAAGGAGGCGGTTGGT  R: CGGGAGTAGCAGTTAGGCTC |
| 8 | 18s^ǂ^ | 64ºC | 175 | F: GGAGAGGGAGCCTGAGAAAC  R: CCTCCAATGGATCCTCGTTA |

^ǂ^Represents primer sequences from our previously published study (18).
