## Supplementary method for "Transient systemic inflammation in adult male mice results in underweight progeny"

**Supplementary methods**

*Behavioural assay*

Anxiety and locomotor activity were assessed using an open field test (1). The test apparatus used in the study consisted of plexiglas arena (transparent walls and floor; W30 × D30 x H30 cm), with the floor divided into nine equal quadrants. Individual animals were gently placed in the center and allowed to freely explore the arena for a preliminary 5 min period. Animal behaviour was then video-recorded for 5 min. The distance travelled, time spent in the center area and numbers of rears was analysed (Ethovision 8.0; Noldus, Wageningen, Netherlands). After each individual test session, the arena was cleaned with 70% alcohol to remove any traces left by the previous animal.

**Reference**

1. Seibenhener ML, Wooten MC. Use of the Open Field Maze to measure locomotor and anxiety-like behavior in mice. J Vis Exp. 2015;(96):e52434.
