## Supplementary figures and images for "Transient systemic inflammation in adult male mice results in underweight progeny"

### Supplementary figure S1

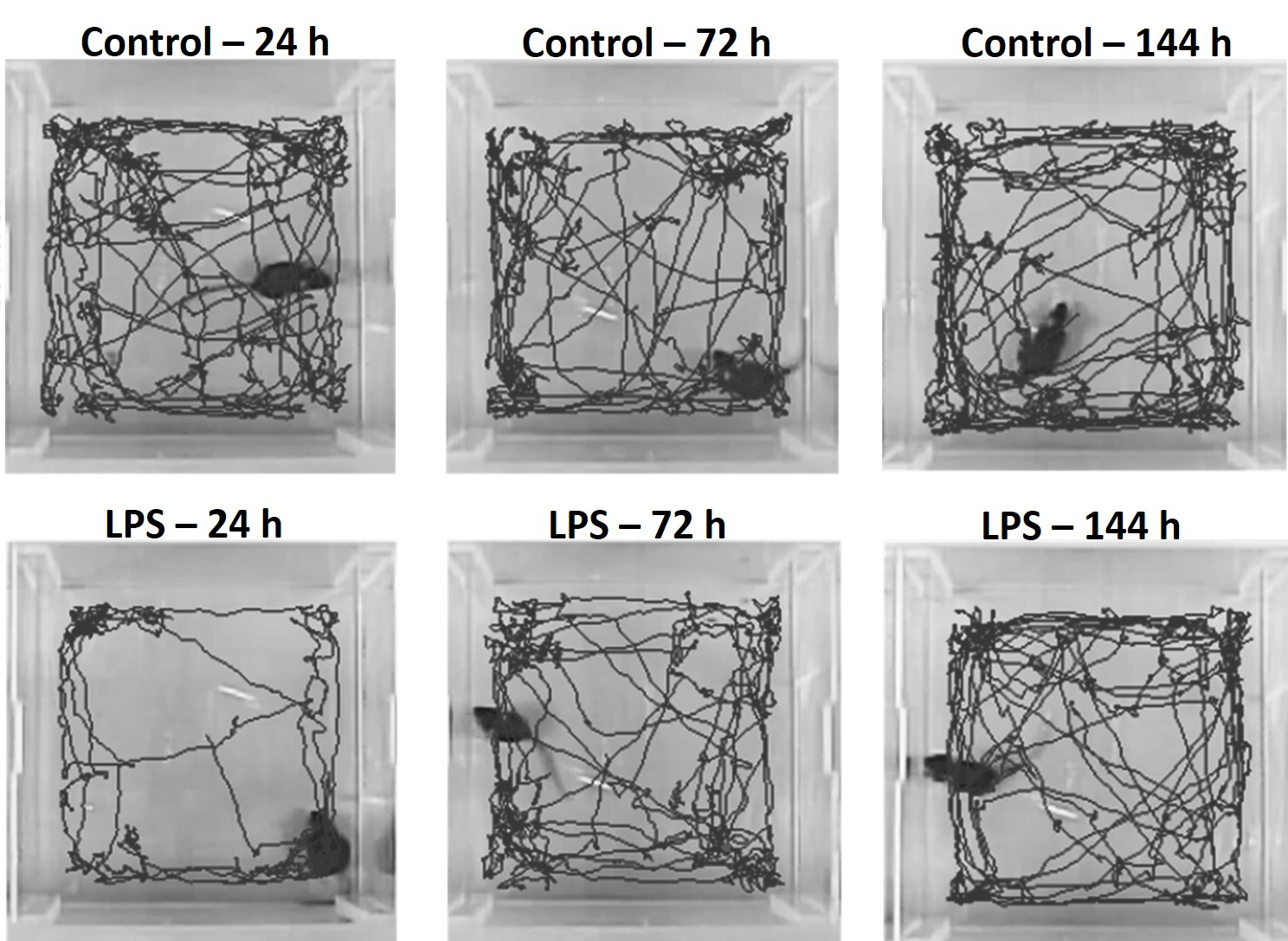

### Supplementary figure S2

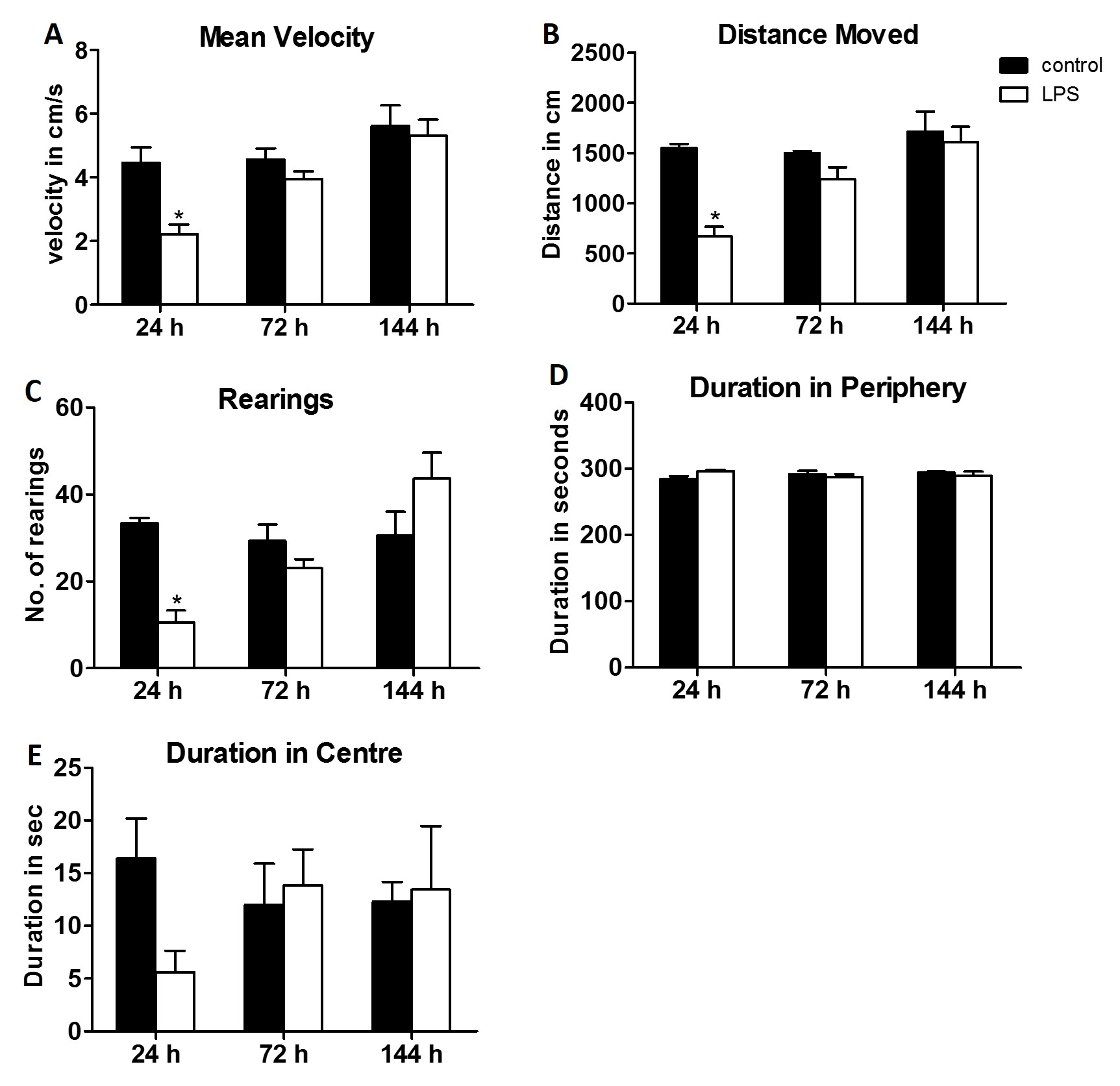

### Supplementary figure S3

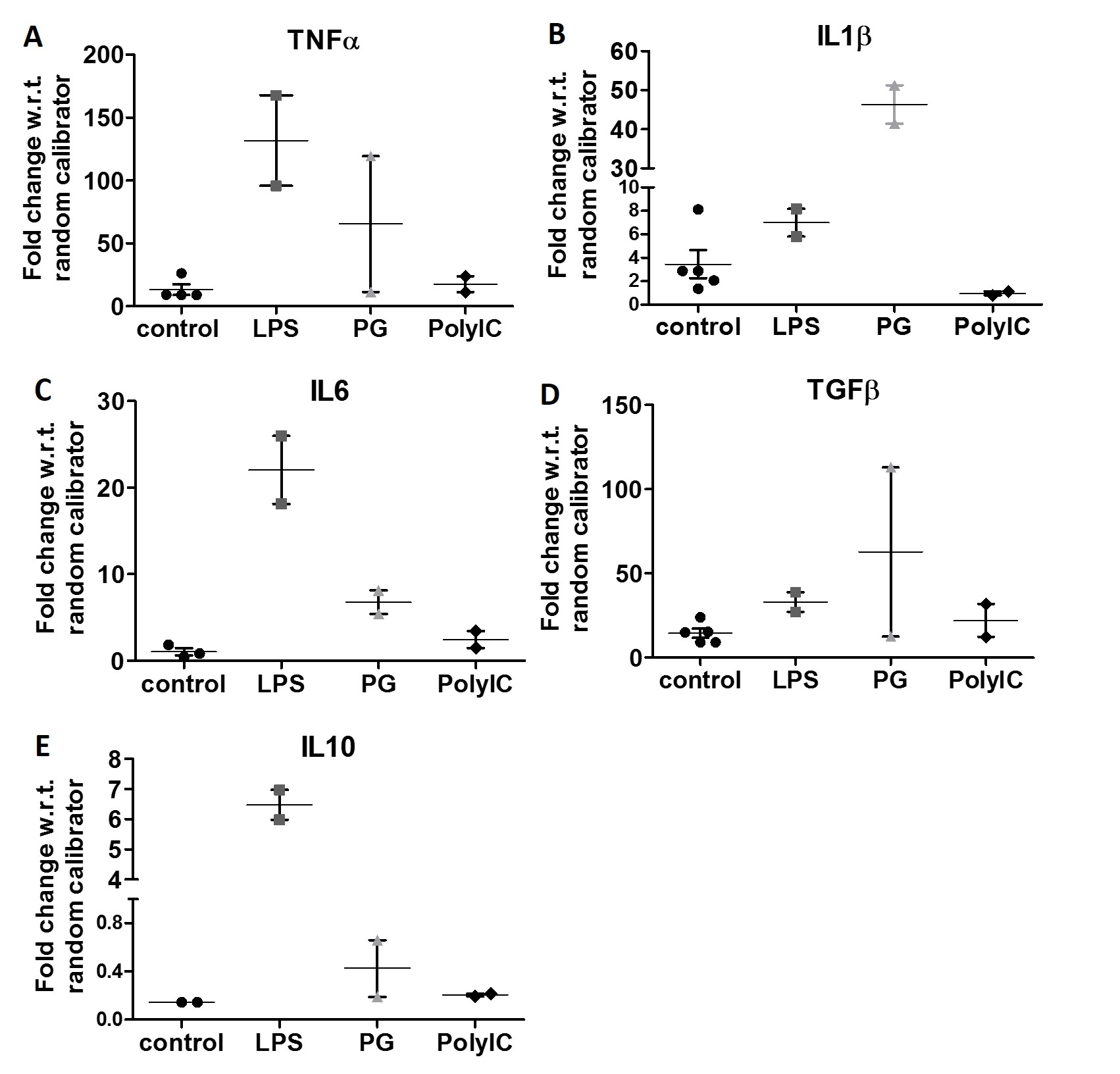

### Supplementary figure S4

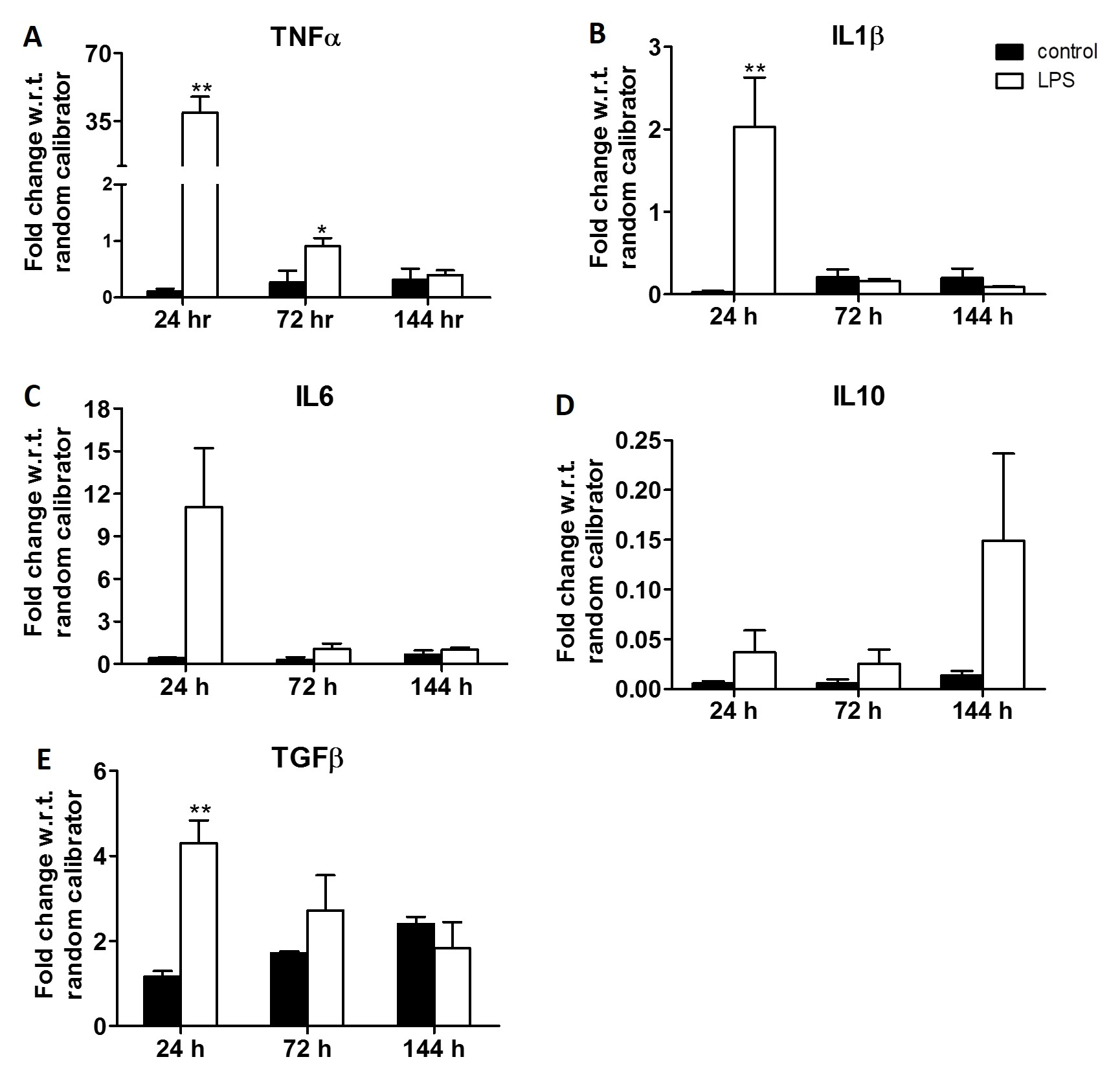

### Supplementary figure S5

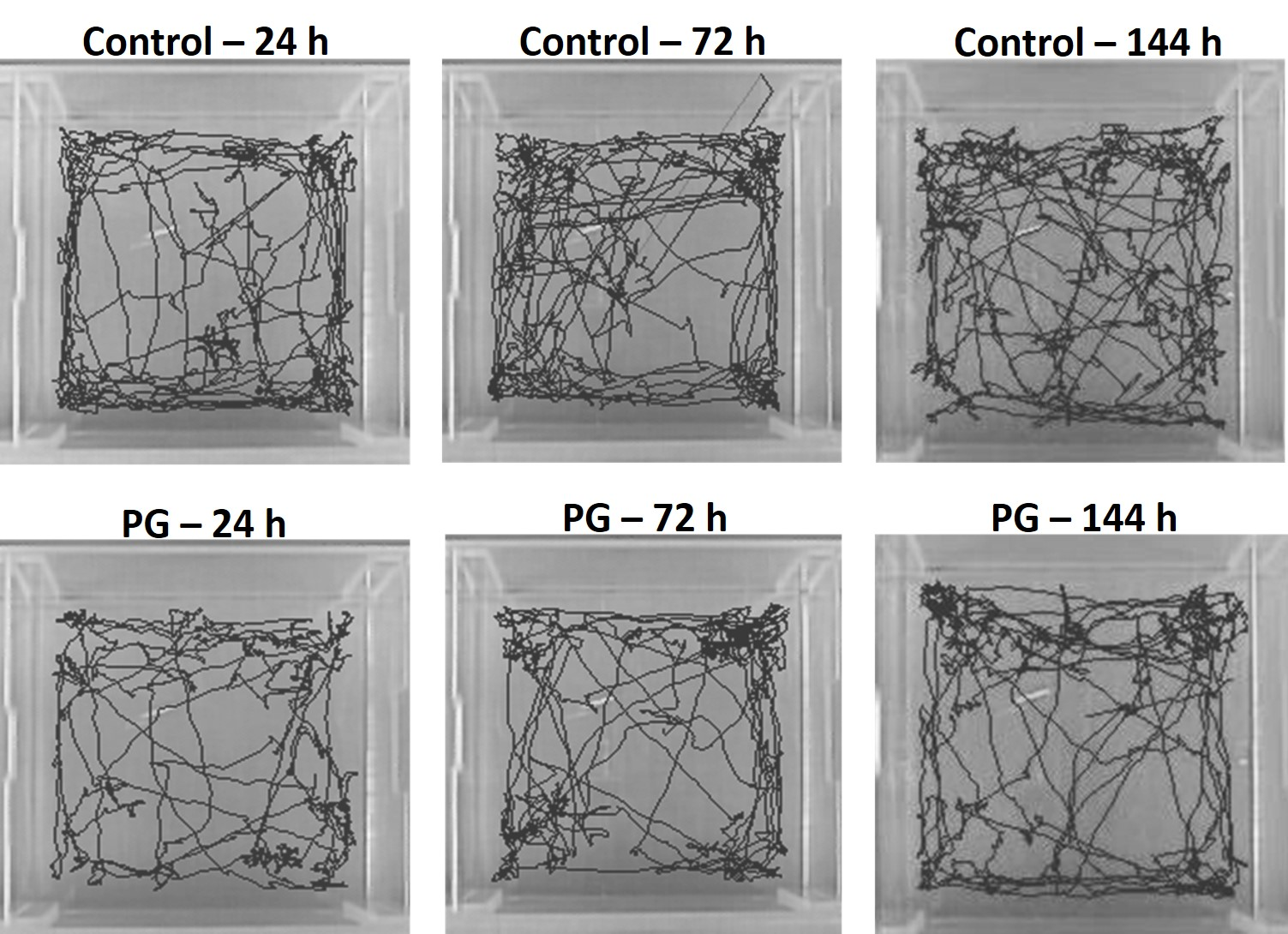

### Supplementary figure S6

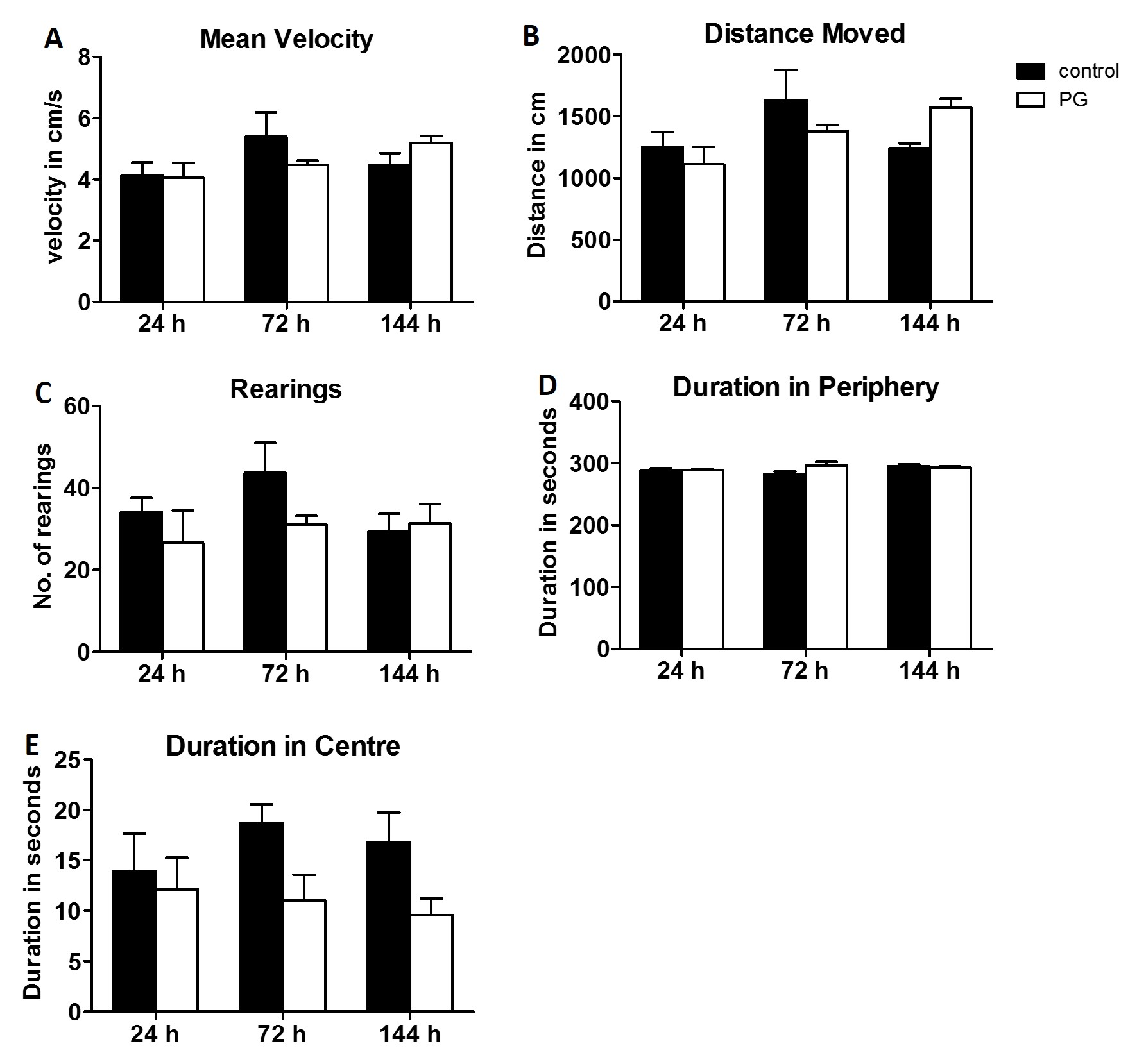

### Supplementary figure S7

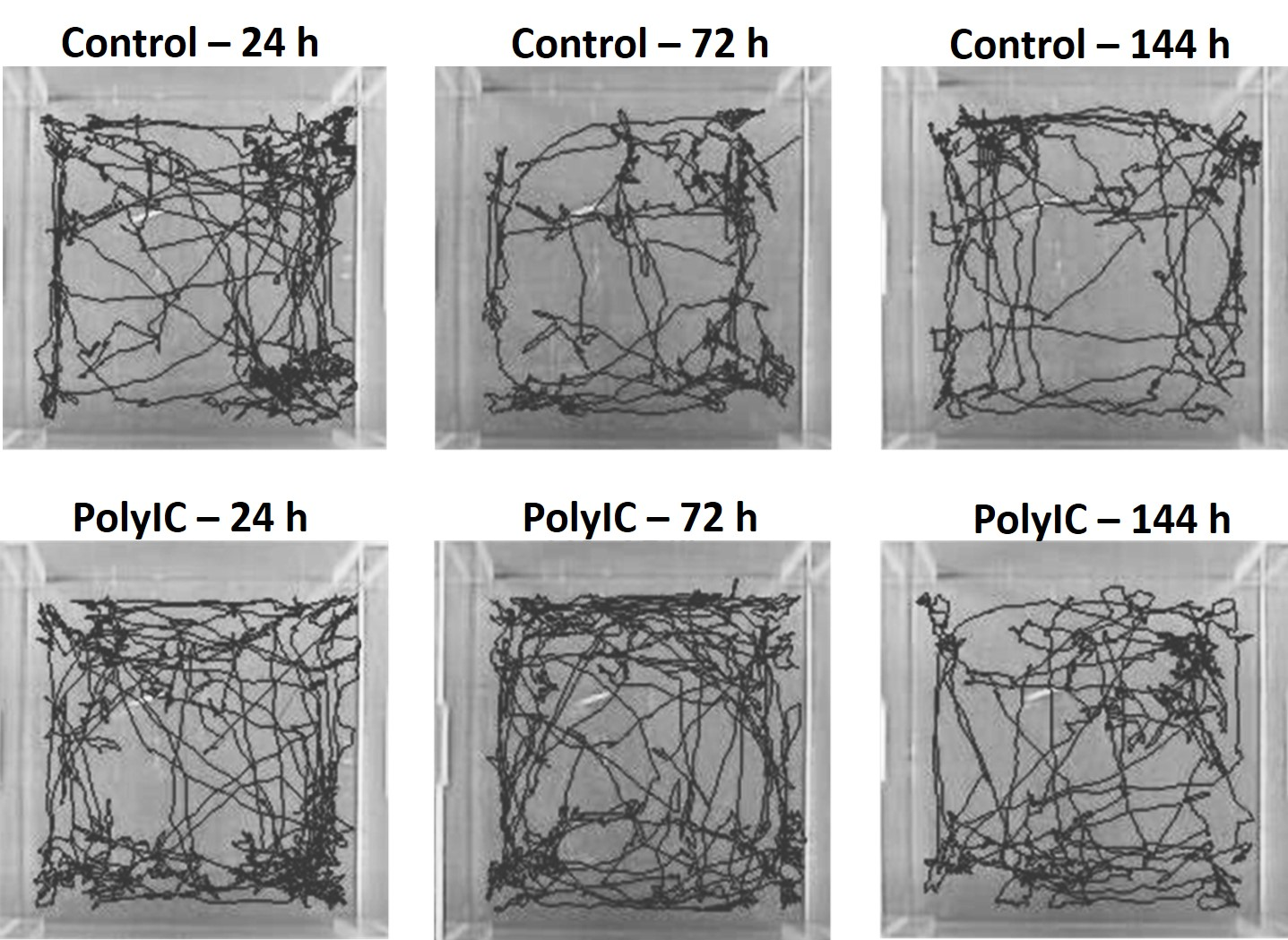

### Supplementary figure S8

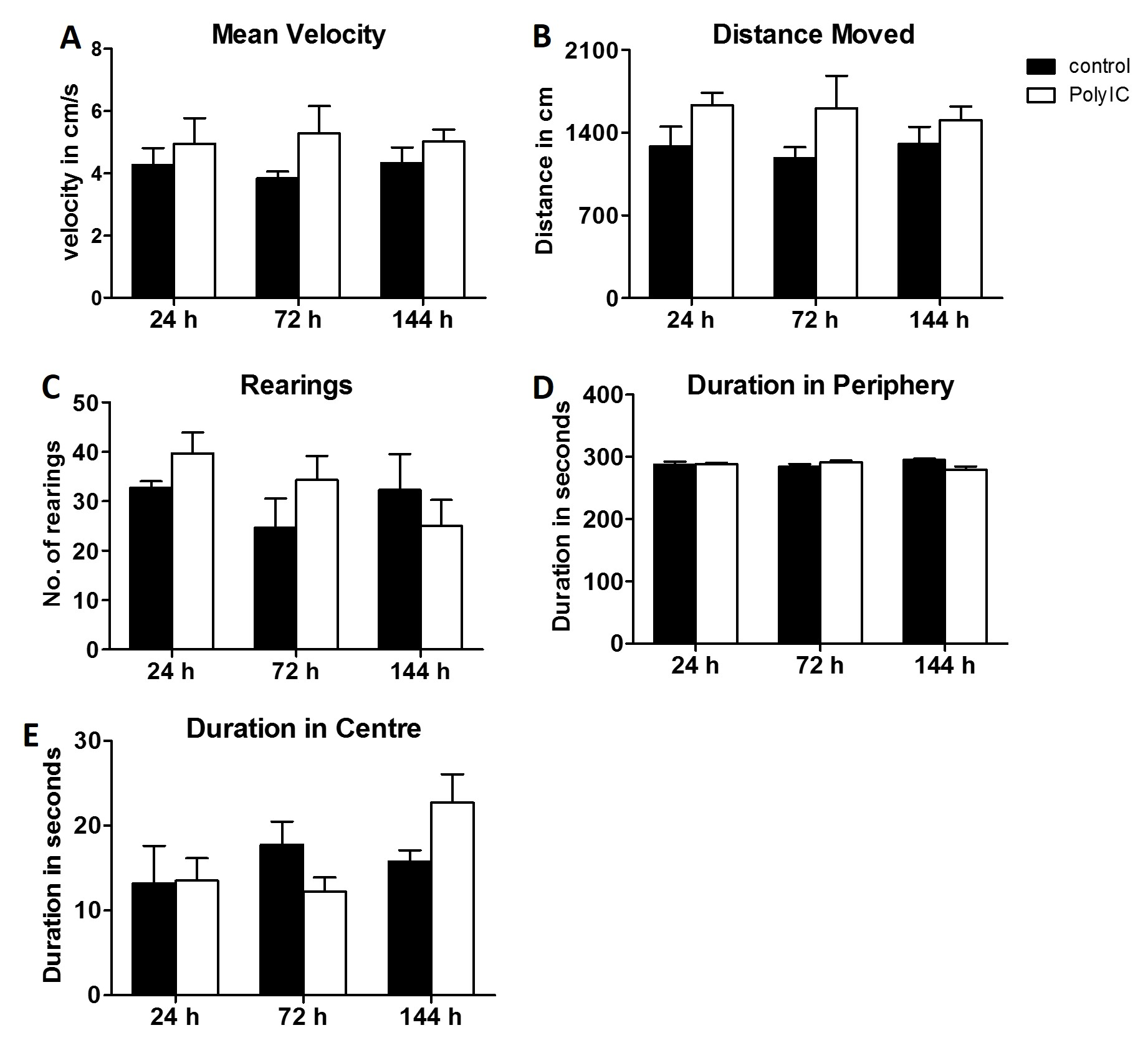

### Supplementary figure S9

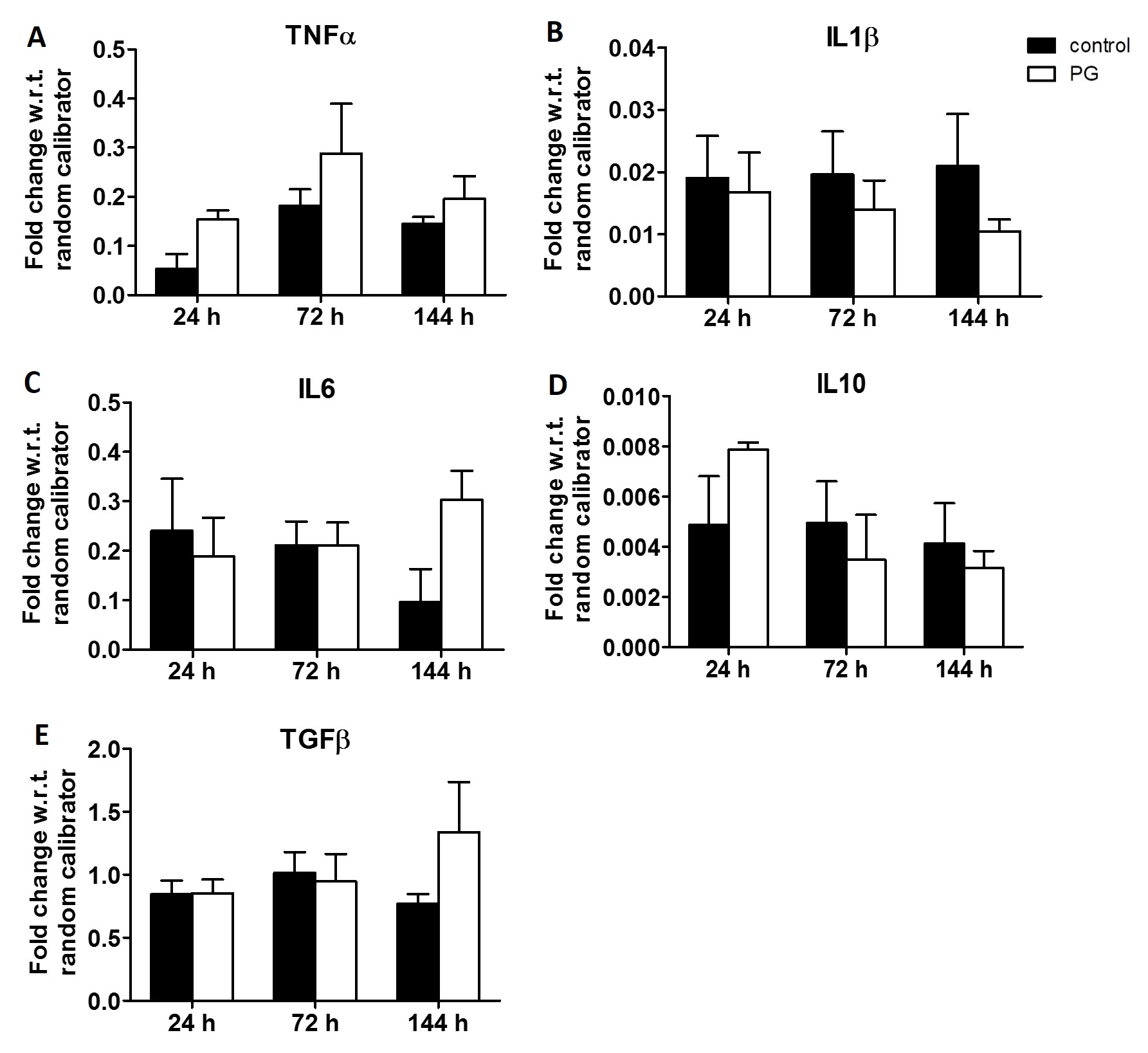

### Supplementary figure S10

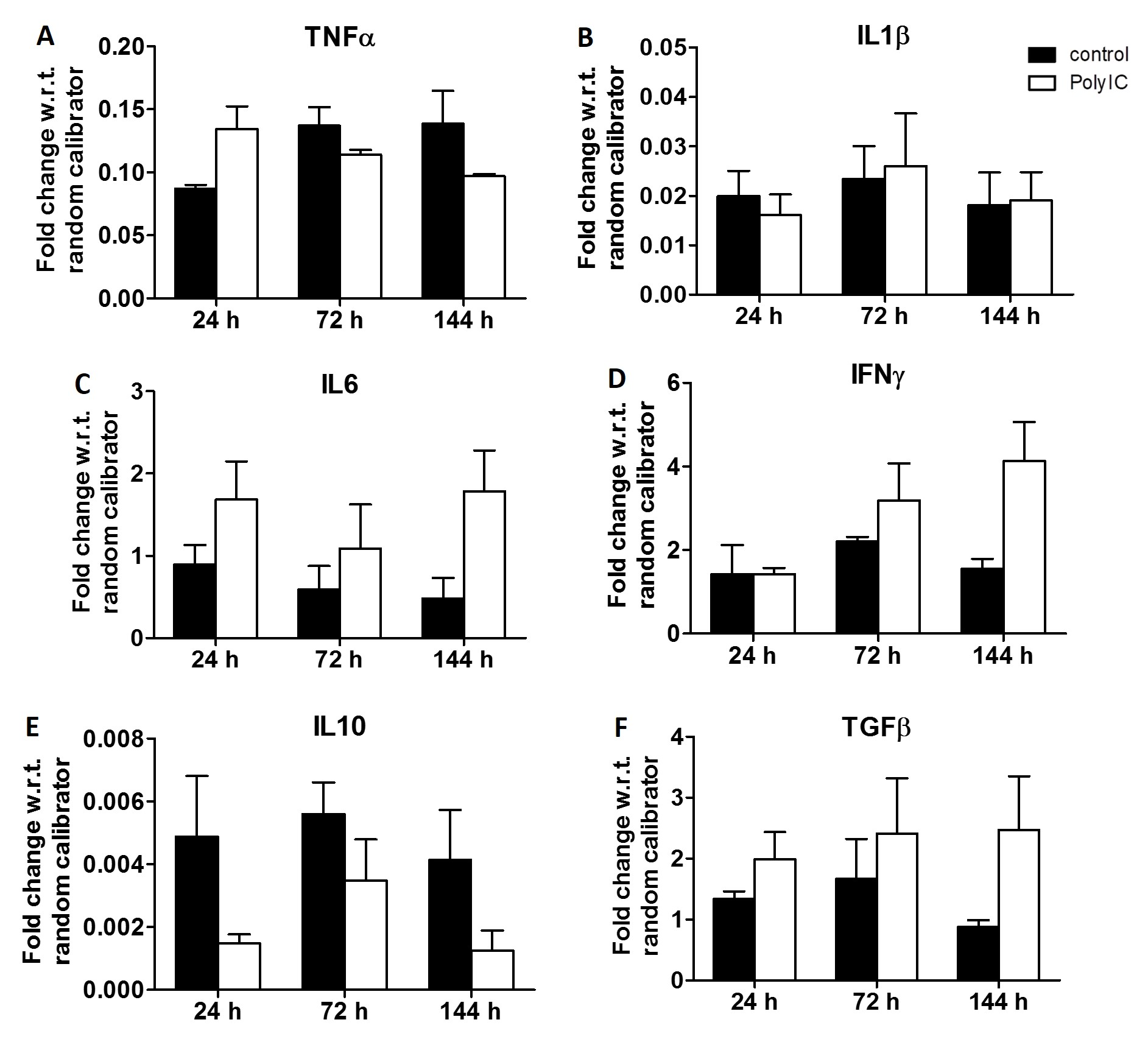
